# Peroxisomal import is circadian in glia and regulates sleep and lipid metabolism

**DOI:** 10.1101/2025.06.23.661129

**Authors:** Anurag Das, Irma Magaly Rivas-Serna, Ankur Kumar, Lakpa Sherpa, Kerui Huang, Hia Kalita, Marlene Dorneich-Hayes, Ruiqi Liu, Vera Mazurak, John P. Vaughen, Hua Bai

**Affiliations:** Department of Genetics, Development, and Cell Biology, Iowa State University, Ames, IA 50011, USA; Interdepartmental Neuroscience PhD program, Iowa State University, Ames, IA 50011, USA; Department of Agriculture, Food, and Nutritional Science, University of Alberta; Edmonton, T6G 2R3, Canada; Interdepartmental Genetics & Genomics PhD program, Iowa State University, Ames, IA 50011, USA; Interdepartmental MCDB PhD program, Iowa State University, Ames, IA 50011, USA; Department of Anatomy, University of California San Francisco, San Francisco, CA 94143, USA

## Abstract

Peroxisomes are critical organelles that detoxify cellular waste while also catabolizing and anabolizing lipids. How peroxisomes coordinate protein import and support metabolic functions across complex tissues and timescales remains poorly understood *in vivo*. Using the *Drosophila* brain, we discover a striking enrichment of peroxisomes in the neuronal soma and the cortex glia that enwrap them. Unexpectedly, import of peroxisomal proteins into cortex glia, but not neurons, oscillated across time and peaked in the early morning. Rhythmic peroxisomal import in cortex glia autonomously required the circadian clock and Peroxin 5 (Pex5; peroxisomal biogenesis factor 5 homolog), with import persistently elevated in clock mutants. Notably, reducing *Pex5* in cortex glia, but not neurons, caused hyperactivity and reduced total sleep. Moreover, brain lipid metabolism was dramatically altered upon *Pex5* knockdown, with glia impacting sphingolipids and triacylglycerols, and neurons impacting phospholipids. The cell-type specificity of these Pex5 phenotypes highlights unique roles for peroxisomal import in both sleep and lipid metabolism in the brain.

## Introduction

Peroxisomes are responsible for vital metabolic activities ranging from lipid metabolism by beta-oxidation and ether lipid biosynthesis[1],[2],[3] to reactive oxygen species detoxification[4],[5],[6]. Accordingly, mutations afflicting peroxisomes can cause numerous diseases, including Zellweger Syndrome[7],[8], a severe peroxisomal biogenesis disorder stemming from mutations in one of the 12 different PEX genes in humans[9]. While these diseases often manifest in the brain[10],[11],[12], why these diseases occur and how peroxisomes are regulated across diverse brain cells and timescales remains incompletely understood.

Peroxisomes require a unique import machinery capable of translocating fully folded and oligomeric substrates from the cytosol into the peroxisomal matrix (lumen)[13]. The Peroxin-5 protein Pex5 is a vital cog in this translocation machinery, as it functions as a major receptor for targeting proteins to the peroxisomal matrix by the peroxisomal targeting signal 1, or PTS1 (e.g. Ser-Lys-Leu, SKL)[14],[15],[16]. Notably, loss of Pex5 from *Drosophila* to mice and humans causes severe defects, including motor and cognitive decline simultaneously with demyelination, axonal loss, and gliosis[17]. While *Pex5* mutants are lethal[18], *Pex5* disruption selectively in mouse oligodendrocytes caused pronounced neurological and lipidomic defects[5]. In contrast, neuronal or astrocytic *Pex5* disruption led to mild defects[17]. While peroxisomal import may thus have cell-type specific effects in the brain, this has not been systematically explored.

*Drosophila* is an outstanding genetic model to interrogate peroxisomes and peroxisomal import function in the brain, given the capability to genetically track and manipulate peroxisomal import in a cell-type specific manner. The fly brain harbors a diversity of glial subtypes and neurons thought to share homologous functions with mammalian counterparts including astrocytes and microglia[19–23]. For example, neuronal synaptic processes are tiled by astrocyte-like glia in mammals[24] and insects[25], and fly axons are isolated and insulated by ensheathing (CNS) and wrapping glia (PNS), analogous to oligodendrocytes and Schwann cells[23,26,27]. In mammals, satellite glia enwrap single neuronal soma[28], while in adult flies, cortex glia encompass dozens of neuronal soma in a honeycombed morphology, and accordingly regulate neuronal activity among other functions[22],[29],[30],[31],[32],[33],[19].

Alongside functionally homologous glia, flies also display complex behavioral repertoires ranging from memory to sleep[34],[35],[36],[37],[38]. While all behavior is tightly coordinated by rhythmic changes experienced under the shared 24-hour light/dark cycle[39],[40], in complex metazoan brains the clock is controlled by a network of circadian pacemaker neurons [41],[42],[43],[44],[45],[46],[47]. Within these cells, negative transcriptional feedback loops between Period (Period/PER), Clock (Clock/CLK), and Cycle (cyc/BMAL1) enforce a ∼24hour molecular rhythm[48]. Glia also express core circadian clock machinery and can modulate sleep through myriad pathways, including remodeling neuronal synapses and regulating metabolism[41,49–51]. Moreover, autonomous disruption of glial clocks directly disrupts sleep and circadian plasticity in clock pacemaker neurons in flies[44]. Although circadian rhythms govern metabolic homeostasis across organisms[52],[53],[54] and in diseases[55], how subcellular organelle dynamics in glia and neurons change across time remains underexplored. Here, we tested how peroxisomal import is coordinated in the fly brain in a cell-type specific manner. We discovered that cortex glia are hubs of abundant matrix peroxisomal proteins and high peroxisomal import, in contrast to other brain glia. Moreover, peroxisomal import in cortex glia but not neurons varied dramatically over circadian time, with elevated import in the early morning. Through genetic analyses, we determined that Pex5 and the core circadian clock autonomously regulates oscillatory peroxisomal import in cortex glia. Notably, loss of Pex5 in cortex glia disrupted sleep behavior and caused ectopic lipid droplets, while loss of Pex5 in all glia or all neurons profoundly altered multiple lipid classes. Together, our work identifies circadian fluctuations in peroxisomal import to specialized glia, which require Pex5 for sleep behavior, and highlights cell-type specific roles of Pex5 in brain lipid metabolism.

## Results

### Peroxisomal abundance and import are enriched in neurons and cortex glia

Peroxisomal import (Figure 1A) is the active translocation of matrix proteins from the cytosol into the peroxisomal lumen via the canonical peroxisomal targeting signal 1 (PTS1; the most common consensus tripeptide sequence is Serine-Lysine-Leucine, SKL). These sequences are recognized by the cytosolic import receptor Pex5, which facilitates shuttling by releasing imported cargo in the peroxisomal lumen,[56]. We first sought to survey peroxisomal import in the adult CNS by co-staining for the peroxisomal marker PMP70 (Peroxisomal membrane protein 70)[57],[16],[58],[59] and *UAS-eYFP.PTS1*[60], which revealed cell-type specific patterns of peroxisomal import when crossed to GAL4 driver lines targeting major CNS populations (Figure 1B). Both pan-glial (*repo-GAL4*) and pan-neuronal (*nSyb-GAL4*) drivers expressed PTS1 puncta that colocalized with PMP70 (Figure 1C). Notably, PMP70 was enriched at the cortex in pan-glial experiments, suggesting that cortex glia may have particularly high levels of peroxisomal import. Indeed, by genetically tracking peroxisomal import with *UAS-eYFP.PTS1* across different glial subtypes, we found that PTS1+,PMP70+ puncta were most abundant in cortex glia compared to other glial classes (Figure 1D-F; Figure S1A-D). Thus, among the CNS glia, cortex glia have uniquely heightened peroxisomal import.

**Figure 1.**
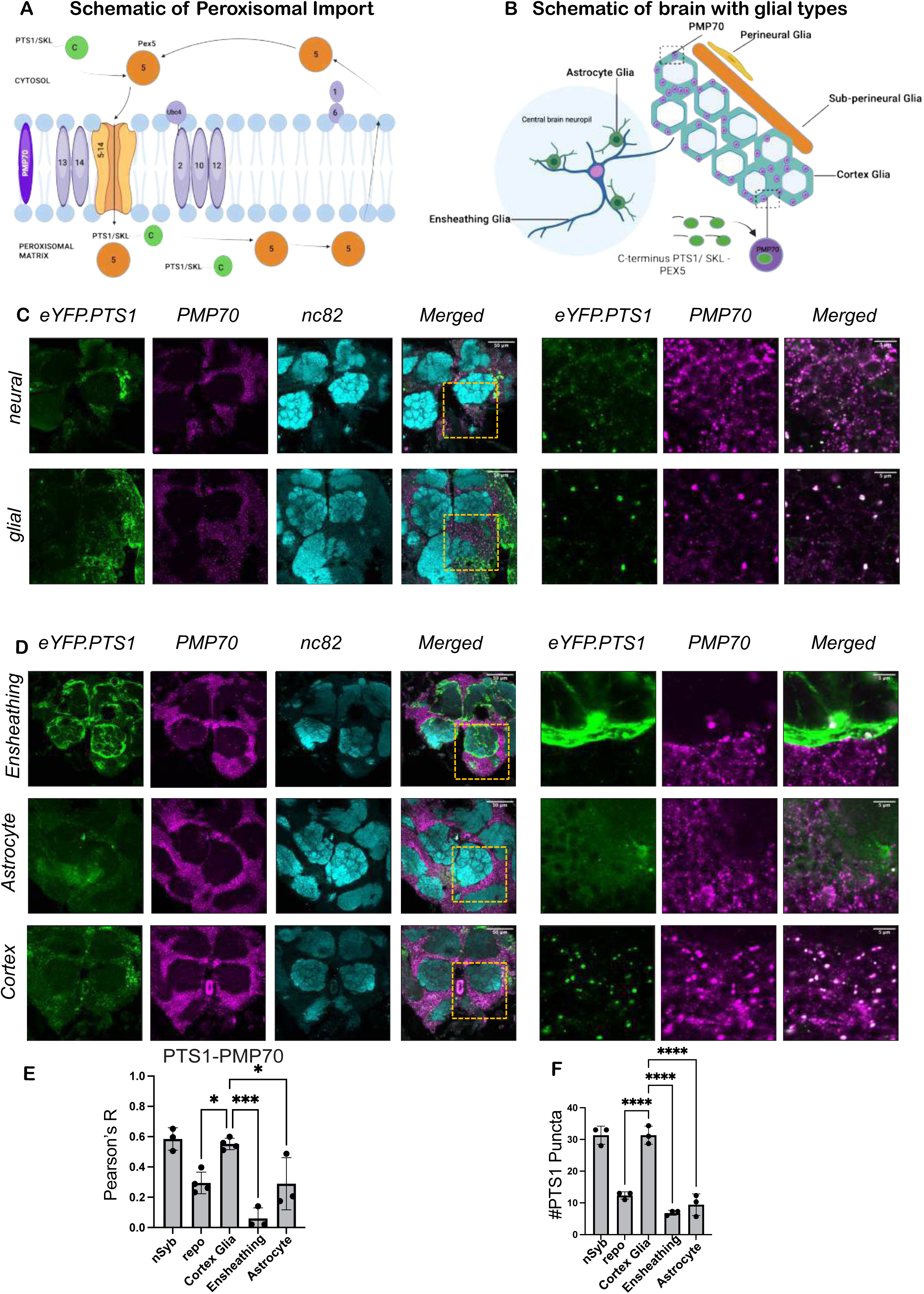
Peroxisomal abundance and import are enriched in neurons and cortex glia. **(A)** Schema of peroxisomal import machinery mediated by Pex5, the cytosolic import receptor that recognizes a *PTS1-SKL* harboring matrix protein and permits its active translocation from the cytosol to the peroxisomal lumen; Adapted from [15] and,[56]. **(B)** Schematic of different glial cell types in the *Drosophila* central nervous system. **(C)** Adult female *Drosophila* brains (7 days old) tracking peroxisome import in neurons with *UAS-eYFP.PTS1* using the pan-neural driver *nSyb-GAL4* or the pan-glial driver *repo- GAL4*. eYFP-tagged *PTS1*-protein import (green) is counterstained with PMP70 (magenta), an abundant peroxisomal membrane protein, and the pre-synaptic active zone protein bruchpilot (cyan; nc82). Yellow rectangle denotes area used for zooms into cortex below the antennal lobes and above the suboesophageal ganglion. Scale bars = 50μm and 5μm (zoom). **(D)** Adult female *Drosophila* brains (7 days old) tracking peroxisomal import in glia subtypes using *UAS-eYFP.PTS1* (green) driven by *GMR56F03-*GAL4 (Ensheathing glia), *GMR83E10-GAL4* (Astrocyte-like glia), and cortex glia (*GMR54H02-*GAL4), counterstained with PMP70 (magenta) and nc82 (cyan). Yellow rectangle denotes area used for zooms into cortex below the antennal lobes and above the suboesophageal ganglion. Scale bars = 50μm and 5μm (zoom). **(E)** Quantification of PTS1-YFP and PMP70 Pearson colocalization across different cellular populations. *n* ≥*3* brains per genotype, data points represent biological replicates averaged over ≥*3* ROIs. **(F)** Quantification of the number of PTS1-YFP punctate across different cellular populations. *n* ≥*3* brains per genotype, dots represent biological replicates averaged over ≥*3* ROIs. * p < 0.05, ** p < 0.01, *** p < 0.001, **** p < 0.0001 by Tukey multiple comparisons test.

### Peroxisomal import oscillates in cortex glia across time and requires Pex5

While further characterizing peroxisomal import, we made the serendipitous finding that import fluctuated across circadian time in cortex glia but not neurons (Figure 2A-B; Figure S2I-K; Zeitgeber 0 (ZT0) denotes lights-on). While neurons had consistently high levels of PTS1 that colocalized with PMP70, cortex glia had three-fold higher levels of PTS1 at ZT2 and ZT4 (Figure S2I-K) compared to a nadir at ZT8 (Figure 2C-F; Pearson decreases from 0.6 to 0.2, while PTS1 puncta number decreases from 28 to 8). A similar enrichment of morning import was observed with an anti-SKL antibody (Figure S2B-D). Import oscillations also occurred under constant darkness implicating endogenous clock control (Figure 2G-J; Figure S2L-N). While cortex glia PTS1 puncta remained low in the evening, the colocalization between PTS1 and PMP70 at ZT14 was higher than at ZT8 (Figure 2C), and we often observed diffuse PTS1 membranous signals at ZT14 (Figure S2E).

**Figure 2.**
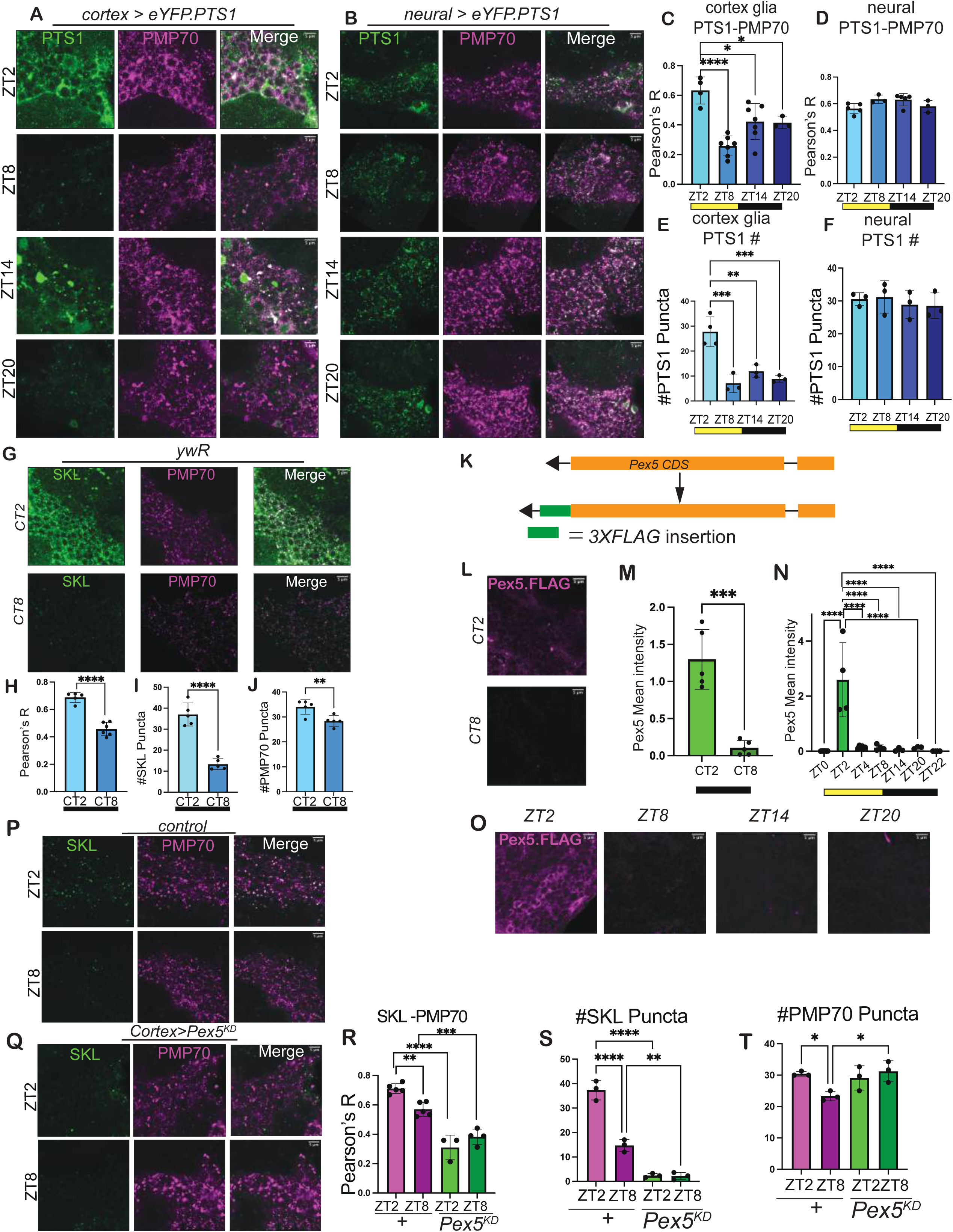
Peroxisomal import oscillates in cortex glia across time and requires Pex5. **(A)** Adult female *Drosophila* brains (7 days old) tracking peroxisomal import oscillation across ZT2, ZT8, ZT14, and ZT20 in cortex glia with *UAS-eYFP.PTS1* crossed to *GMR54H02-GAL4*. eYFP-tagged *PTS1*-protein import (green) is counterstained with PMP70 (magenta), an abundant peroxisomal membrane protein. Scale bars = 5μm. **(B)** Adult female *Drosophila* brains (7 days old) tracking peroxisomal import oscillation across ZT2, ZT8, ZT14, and ZT20 in neurons with *UAS-eYFP.PTS1* crossed to the pan-neural driver *nSyb-GAL4.* eYFP-tagged *PTS1*-protein import (green) is counterstained with PMP70 (magenta), an abundant peroxisomal membrane protein. Scale bars = 5μm. **(C)** Quantification of PTS1-YFP and PMP70 Pearson colocalization across cortex glia. *N* ≥*3* brains per genotype, dots represent biological replicates. ROIs were taken from zooms into cortex below the antennal lobes and above the suboesophageal ganglion. **(D)** Quantification of PTS1-YFP and PMP70 Pearson colocalization across neurons. *N* ≥*3* brains per genotype, dots represent biological replicates. **(E)** Quantification of the number of PTS1-YFP punctate across cortex glia. *n* ≥*3* brains per genotype, dots represent biological replicates averaged over ≥*3* ROIs. **(F)** Quantification of the number of PTS1-YFP punctate across neurons. *n* ≥*3* brains per genotype, dots represent biological replicates averaged over ≥*3* ROIs. **(G)** Adult female *Drosophila* brains (7 days old) *yw^R^* tracking peroxisomal import oscillation across constant darkness comparing CT2 versus CT8 stained with SKL (green) and PMP70 (magenta), an abundant peroxisomal membrane protein. Scale bars = 5μm. **(H)** Quantification of SKL and PMP70 Pearson colocalization. *n* ≥*3* brains per genotype, data points represent biological replicates. ROIs were taken from zooms into cortex below the antennal lobes and above the suboesophageal ganglion. **(I)** Quantification of the number of SKL punctate. *n* ≥*3* brains per genotype, dots represent biological replicates averaged over ≥*3* ROIs. **(J)** Quantification of the number of PMP70 punctate. *n* ≥*3* brains per genotype, dots represent biological replicates averaged over ≥*3* ROIs. **(K)** Schema of *Pex5.FLAG* knock-in generated using CRISPR/Cas9-mediated gene editing in *Drosophila*. **(L)** Adult female *Drosophila* brains (7 days old) tracking *Pex5.FLAG* knock-in stained with *anti-FLAG* (magenta) across CT2, CT8 (CT = circadian time; animals were entrained on light-dark cycles and then kept in constant darkness for one day) **(M)** Quantification of Pex5 mean intensity in constant darkness. *n* ≥*3* brains per genotype, dots represent biological replicates. **(N)** Quantification of Pex5 mean intensity in 12hr light/dark cycle. *n* ≥*3* brains per genotype, dots represent biological replicates. **(O)** Adult female *Drosophila* brains (7 days old) tracking *Pex5.FLAG* knock-in stained with *anti-FLAG* (magenta) across ZT2, ZT8, ZT14, and ZT20. **(P)** Adult female *Drosophila* brains (7 days old) stained with SKL (green) and PMP70 (magenta) in control *w^1118^/UAS-Pex5 RNAi*. **(Q)** Adult female *Drosophila* brains (7 days old) stained with SKL (green) and PMP70 (magenta) in cortex-glia specific knockdown of Pex5 using *GMR54H02-Gal4*> *UAS-Pex5 RNAi.* ROIs were taken from zooms into cortex below the antennal lobes and above the suboesophageal ganglion. **(R)** Quantification of SKL and PMP70 Pearson colocalization. *n* ≥*3* brains per genotype, data points represent biological replicates. **(S)** Quantification of the number of SKL punctate. *n* ≥*3* brains per genotype, dots represent biological replicates averaged over ≥*3* ROIs. **(T)** Quantification of the number of PMP70 punctate. *n* ≥*3* brains per genotype, dots represent biological replicates averaged over ≥*3* ROIs. * p < 0.05, ** p < 0.01, *** p < 0.001, **** p < 0.0001 by Tukey’s multiple comparisons test.

Intrigued, we tested whether the central regulator of peroxisomal import, Pex5 (Figure 1A), also oscillated in the brain. We used CRISPR to generate a knock-in FLAG tag into the endogenous Pex5 locus (Figure 2K; Figure S2G and see Methods). Importantly, *Pex5.FLAG* flies were viable and had wild type levels of Pex5 transcript (Figure S2F-H’), unlike Pex5 loss-of-function alleles, which are lethal[18]. Using *Pex5.*FLAG, we observed that Pex5 protein robustly oscillated across time in the brain under light-dark and dark-dark conditions, peaking at ZT2 during the height of PTS1 import in cortex glia (Figure 2L-O; Figure S2A; note that the majority of Pex5 is expected to be cytosolic). Moreover, this oscillatory signature is present in single cell RNA sequencing data, where transcripts of *Pex5*, *Pex11*, and the peroxisome proliferator-activated receptor (PPAR) homolog *Eip75B* peak at similar times in cortex glia[43] (Figure S3 A-B). To test if Pex5 is obligatory for basal levels of peroxisomal import in glia, we knocked-down *Pex5* using an *RNAi* that reduces *Pex5* transcripts by over 50% [4]. *Pex5^KD^* specifically in cortex glia strongly suppressed overall peroxisomal import, with reduced PTS1 puncta that no longer oscillated across time, but did not alter PMP70+ puncta (Figure 2P-T). This phenotype was replicated with a second Pex5 *RNAi* line (Figure S4A-D). Thus, in cortex glia, peroxisomal import and Pex5 levels are highest in the morning, and oscillations in peroxisomal import require Pex5.

### The circadian clock autonomously controls peroxisomal import in cortex glia

To test if the circadian clock regulated peroxisomal import, we utilized *period* arrhythmic mutants (*per^01^*), and measured PTS1 import in cortex glia at the apex (ZT2) and nadir (ZT8) of import. Strikingly, *per* mutants had persistently elevated PTS1 import in cortex glia (Figure 3A-E). As all glia and subsets of neurons express the core circadian clock machinery[61],[62],[63],[64],[65],[43], we next investigated whether cortex glia autonomously require a functioning circadian clock by leveraging a dominant-negative Clock transgene[66] (*UAS-Clk^DN^*) or a validated *per RNAi line* [67],[68,69]. In both manipulations, autonomously disrupting the cortex glia circadian clock triggered aberrant and persistently elevated levels of PMP70/SKL at ZT8 (Figure 3F-J; Figure S6A-E). Additionally, removing the PPAR homolog and downstream target of Per, *Eip75B*, in cortex glia phenocopied the effects of *per* and *Clk^DN^*and persistently elevated import (Figure S5A-C). Thus, the circadian clock autonomously controls rhythmic peroxisomal import into a specialized subset of brain cells, the cortex glia.

**Figure 3.**
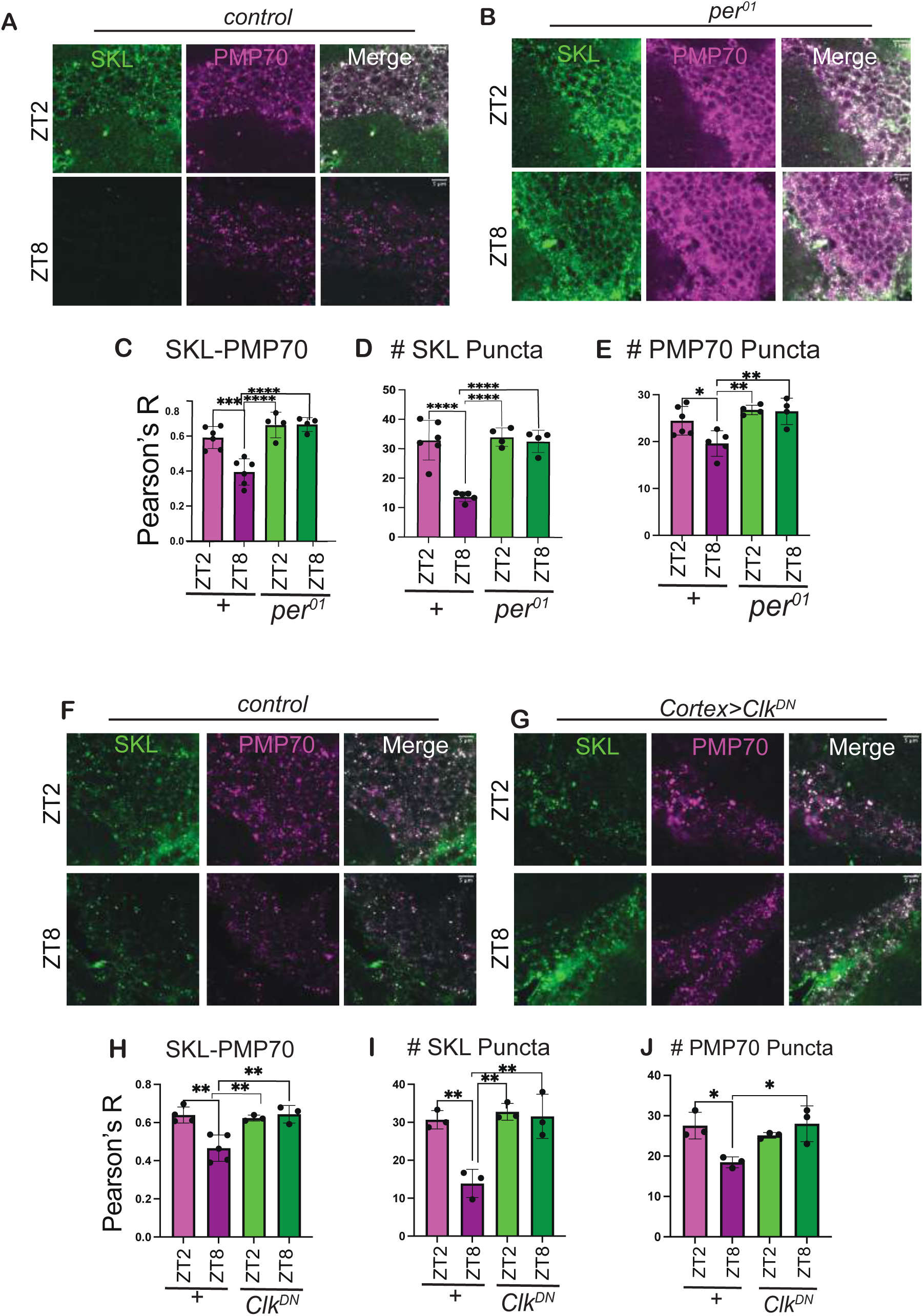
The circadian clock autonomously controls peroxisomal import in cortex glia. **(A)** Adult female *Drosophila* brains (7 days old) tracking peroxisomal import oscillation across ZT2, ZT8 in Control (*CantonS*) with SKL (green) counterstained with PMP70 (magenta). **(B)** Adult female *Drosophila* brains (7 days old) tracking peroxisomal import oscillation across ZT2, ZT8 in the clock mutant *Per^01^* with SKL (green) counterstained with PMP70 (magenta). **(C)** Quantification of SKL and PMP70 Pearson colocalization. *n* ≥*3* brains per genotype, data points represent biological replicates averaged over ≥*3* ROIs. ROIs were taken from zooms into cortex below the antennal lobes and above the suboesophageal ganglion. **(D)** Quantification of the number of SKL punctate. *n* ≥*3* brains per genotype, dots represent biological replicates averaged over ≥*3* ROIs. **(E)** Quantification of the number of PMP70 punctate. *n* ≥*3* brains per genotype, dots represent biological replicates averaged over ≥*3* ROIs. **(F)** Adult female *Drosophila* brains (7 days old) stained with SKL (green) and PMP70 (magenta) in UAS-control (*w^1118^/UAS-CLK-DN*) **(G)** Adult female *Drosophila* brains (7 days old) stained with SKL (green) and PMP70 (magenta) in Cortex-Glia>*UAS-Clk-Dominant negative(DN)* – *GMR54H04-Gal4>UAS-CLK-DN*. **(H)** Quantification of SKL and PMP70 Pearson colocalization. *n* ≥*3* brains per genotype, data points represent biological replicates averaged over ≥*3* ROIs. ROIs were taken from zooms into cortex below the antennal lobes and above the suboesophageal ganglion. **(I)** Quantification of the number of SKL punctate. *n* ≥*3* brains per genotype, dots represent biological replicates averaged over ≥*3* ROIs. **(J)** Quantification of the number of PMP70 punctate. *n* ≥*3* brains per genotype, dots represent biological replicates averaged over ≥*3* ROIs. ns – non-significant, * p < 0.05, ** p < 0.01, *** p < 0.001, **** p < 0.0001 by Tukey’s multiple comparisons test.

### Loss of *Pex5* selectively in cortex glia disrupts sleep

Given the striking circadian oscillations in peroxisomal import in cortex glia, and that cortex glia regulate sleep behavior[70],[71], we next examined whether removing *Pex5* in different cells altered sleep. Remarkably, disrupting *Pex5* in cortex glia reduced both daytime and nighttime sleep, with shorter sleep bout duration (Figure 4A-D). This phenotype likely stems from hyperactivity, as total daytime activity was elevated in *Pex5^KD^* flies (Figure 4E-G). Similar deficits occurred under constant darkness (Figure 4H-N), and there were no clear defects in sleep latency or circadian rhythmicity (Figure S7A-B). In contrast, knocking-down *Pex5* in ensheathing glia had no impact on daytime sleep and modest effects on nighttime sleep (Figure S7K-R). Similarly, removing *Pex5* from neurons with *nSyb-GAL4* did not consistently affect daytime or nighttime sleep (Figure S7C-J). Thus, Pex5 plays a critical, cell-type specific role in cortex glia that regulates activity and total sleep.

**Figure 4.**
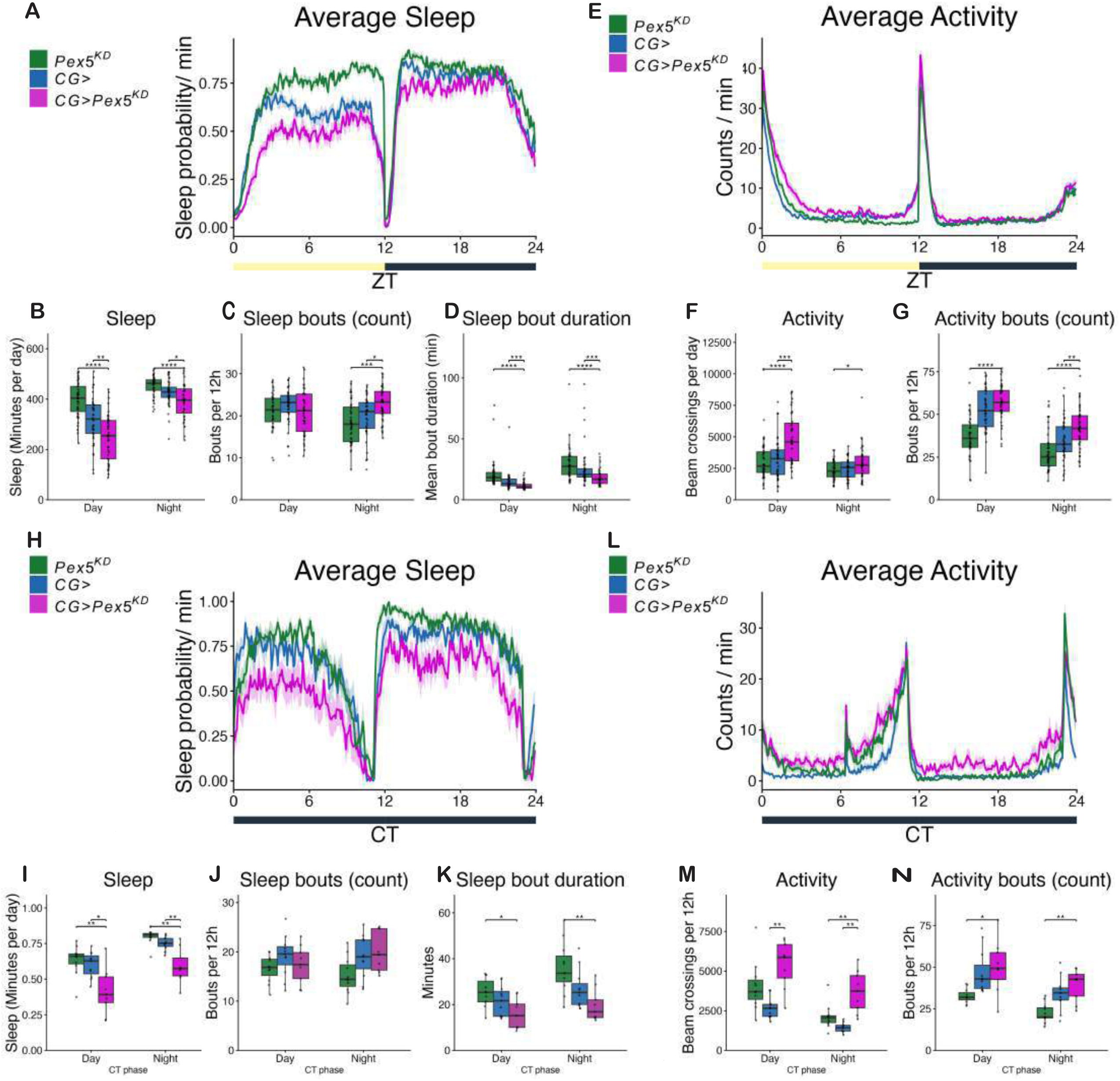
Loss of *Pex5* selectively in cortex glia disrupts sleep. **(A)** Average Sleep (sleep probability/min) was measured for 12hr/12hr, light/dark cycle on an average for 4-5 days continuously for 5-day to 10-day old males post-eclosion. Average sleep time-plot showing sleep probability/min (bouts of inactivity > 5 mins) for the cortex glia driver *GMR77A03-GAL4* crossed with *UAS-Pex5 RNAi-KD* (magenta) compared to parental controls *UAS-Pex5-RNAi (*green) *& GMR77A03-GAL4 (blue)*. **(B)** Total sleep(minutes per day) quantified for all three genotypes from **(A).** Individual black dots represent different biological replicates. **(C)** Sleep bouts per 12hour quantified for all three genotypes from **(A).** Individual black dots represent different biological replicates. **(D)** Sleep bout duration per minute quantified for all three genotypes from **(A).** Individual black dots represent different biological replicates. **(E)** Average activity was measured for 12hr/12hr, light/dark cycle on an average for 4-5 days continuously for 5-day to 10-day old males post-eclosion. Average activity time-plot showing counts/min (bouts of inactivity > 5 mins) for the cortex glia driver *GMR77A03- GAL4* crossed with *UAS-Pex5 RNAi KD* (magenta) compared to parental controls *UAS- Pex5-RNAi (*green) *& GMR77A03-GAL4 (blue)*. **(F)** Total activity (beam crossings per day) quantified for all three genotypes from **(E).** Individual black dots represent different biological replicates. **(G)** Activity bouts (per 12 hour) quantified for all three genotypes from **(E).** Individual black dots represent different biological replicates. **(H)** Average Sleep (sleep probability/min) was measured for 12hr/12hr, dark/dark cycle on an average for 5-6 days continuously for 5-day to 10-day old males post-eclosion after initial entrainment on light/dark cycle. Average sleep time-plot showing sleep probability/min (bouts of inactivity > 5 mins) for the cortex glia driver *GMR77A03-GAL4* crossed with *UAS-Pex5 RNAi KD* (magenta) compared to parental controls *UAS-Pex5-RNAi (*green) *& GMR77A03-GAL4 (blue)*. **(I)** Total sleep(minutes per day) quantified for all three genotypes from **(H).** Individual black dots represent different biological replicates. **(J)** Sleep bouts per 12hour quantified for all three genotypes from **(H).** Individual black dots represent different biological replicates. **(K)** Sleep bout duration per minute quantified for all three genotypes from **(H).** Individual black dots represent different biological replicates. **(L)** Average activity was measured for 12hr/12hr, dark/dark cycle on an average for 5-6 days continuously for 5-day to 10-day old males post-eclosion after initial entrainment on light/dark cycle. Average activity time-plot showing counts/min (bouts of inactivity > 5 mins) for the cortex glia driver *GMR77A03-GAL4* crossed with *UAS-Pex5 RNAi KD* (magenta) compared to parental controls *UAS-Pex5-RNAi (*green) *& GMR77A03-GAL4 (blue)*. **(M)** Total activity (beam crossings per day) quantified for all three genotypes from **(L).** Individual black dots represent different biological replicates. **(N)** Activity bouts (per 12 hour) quantified for all three genotypes from **(L).** Individual black dots represent different biological replicates. * p < 0.05, ** p < 0.01, *** p < 0.001, **** p < 0.0001 by ANOVA, Tukey’s multiple comparisons or Welch’s ANOVA depending on equality of data variance. For nonparametric sleep bouts, Mann-Whitney U test was used. n ≥ 40 flies for light/dark experiment (Figure 5A-E); n ≥ 10 flies for dark/dark experiment (Figure 5H-N);

### Pex5 is essential for lipid homeostasis in the adult brain

Intriguingly, wakefulness causes accumulation of lipid droplets in cortex glia and ensheathing glia[71], and peroxisomes and peroxins[72] can directly interact with lipid droplets (LD), with Pex5 stimulating lipolysis[73]. We thus hypothesized that removing *Pex5* would alter lipid metabolism in the brain. Indeed, knockdown of *Pex5* in cortex glia dramatically increased lipid droplet number and size (Figure 5A-C) with lipid droplets appearing adjacent to PMP70 puncta. Knocking down Pex5 in neurons moderately increased lipid droplet number and size (Figure S8J-L). These data are consistent with Pex5 function impacting lipid droplet biology, either through peroxisomal import or direct effects on lipolysis through lipase recruitment [73].

**Figure 5.**
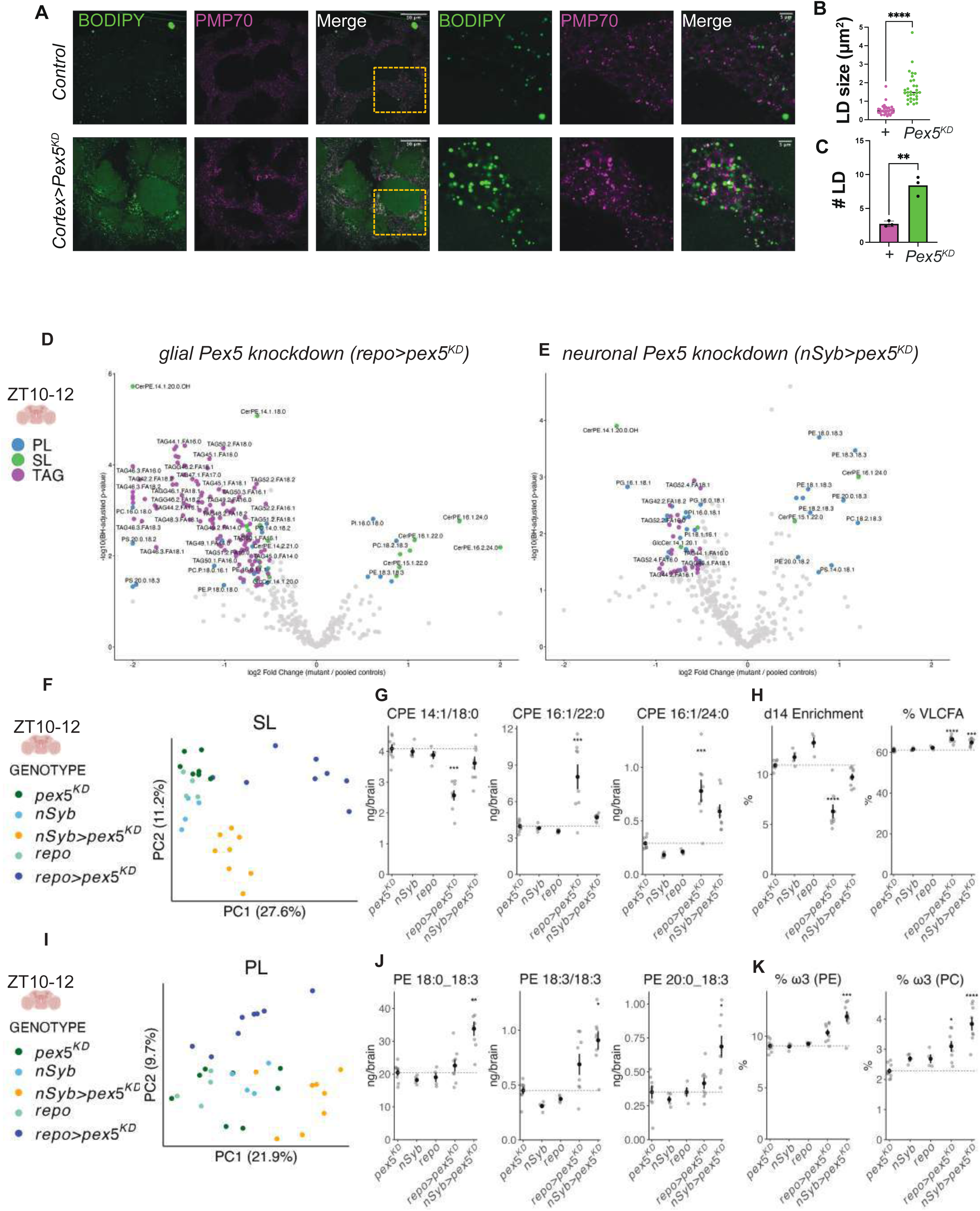
Pex5 is essential for lipid homeostasis in the adult brain. **A)** Adult female *Drosophila* brains (7 days old) tracking neutral lipid droplet (BODIPY 493/503, green) counterstained with PMP70 (magenta). Representative images of UAS-control (*UAS-Pex5 RNAi>w^1118^*) versus cortex-glia specific knockdown of Pex5 using *GMR54H02-Gal4*>*UAS-Pex5 RNAi*. **(B)** Quantification of different lipid droplet size(um^2^) from A. *n* ≥*3* brains per genotype, dots represent individual lipid droplets per biological replicates averaged over ≥*3* ROIs. ROIs were taken from zooms into cortex below the antennal lobes and above the suboesophageal ganglion, and quantified from 200×200 pixel windows inside these zooms. **(C)** Quantification of lipid droplet count from A. *n* ≥*3* brains per genotype, dots represent number of lipid droplets per biological replicates averaged over ≥*3* ROIs. ROIs were taken from zooms into cortex below the antennal lobes and above the suboesophageal ganglion, and quantified from 200×200 pixel windows inside these zooms. **(D)** LC-MS/MS analysis of day 7 female brains at ZT10-12 for glial *Pex5* knockdown; see Table S1 for full genotypic details. Volcano plot shows p-value versus log2foldchange for panglial *pex5^KD^* versus pooled parental controls. Phospholipids are colored blue, sphingolipids colored green, and TAG colored purple. *n* ≥*4* tubes of 20 brains for each genotype. **(E)** LC-MS/MS analysis of day 7 female brains at ZT10-12 for neuronal *Pex5* knockdown. Volcano plot shows p-value versus log2foldchange for pan-neuronal *pex5^KD^* versus pooled parental controls. Phospholipids are colored blue, sphingolipids colored green, and TAG colored purple. **(F)** Principal component analysis of sphingolipids separates glial *pex5* (blue) and neuronal *pex5* knockdowns (orange) from controls. **(F)** Principle component analysis of sphingolipids across the indicated genotypes (orange = neuronal *pex5^KD^*, dark blue = glial *pex5^KD^*, light blue = neuronal *GAL4* control, light green = glial *GAL4* control, dark green = *pex5^KD^* control) **(G)** Total ng/brain for select ceramide phosphoethanolamine (CPE) sphingolipids. **(H)** Sphingolipid metadata analysis, showing the ratio of d14 sphingolipids over total sphingolipids, and the % of very long chain fatty acids (VLCFA; C > 20). **(I)** Principal component analysis of phospholipids separates neuronal *pex5* knockdowns from controls. (orange = neuronal *pex5^KD^*, dark blue = glial *pex5^KD^*, light blue = neuronal *GAL4* control, light green = glial *GAL4* control, dark green = *pex5^KD^* control) **(J)** Example ng/brain values for 18:3(omega-3) fatty acid chain containing species that increase upon neuronal *pex5^KD^*. ω3 fatty acids harbor more than one carbon-carbon double bond, starting from the 3^rd^ carbon away from the terminal of the fatty acid tail. **(K)** % of ω3 lipids in phosphatidylethanolamine (PE) and phosphatidylcholine (PC). * p < 0.05, ** p < 0.01, *** p < 0.001, **** p < 0.0001 by Tukey’s multiple comparisons test.

We next used liquid chromatography-tandem mass spectrometry (LC-MS/MS) for targeted analyses of *Drosophila* sphingolipids (SL), phospholipids (PL; including glycerophospholipids and alky-ether phospholipids); free fatty acids; and triacylglycerols (TAG). We dissected brains from controls (*RNAi* and *GAL4* lines separately), or flies with *Pex5* knocked-down in all glia (*repo-GAL4 > Pex5^KD^*) or all neurons (*nSysb-GAL4 > Pex5^KD^)* at ZT10-12 in day 7 adults. Both neuronal or glial removal of *Pex5* caused dramatic and unique shifts to the global brain lipidome (Figure 5D-E), with more species altered by the glial manipulation, including decreases to individual and total TAGs (Figure 5D; Figure S8A-B). Although aged *Pex5* mutants accumulate very long chain fatty acids (VLCFA) from impaired beta-oxidation, in young day 7 adults we did not detect pronounced VLCFA accumulation in either glial or neuronal *Pex5^KD^*(Figure S8C). Strikingly, glial *Pex5^KD^* significantly and specifically altered major ceramide phosphoethanolamine (CPE) species by decreasing d14 sphingoid bases (d14; long-chain base) and increasing d16 sphingoid bases (Figure 5G). This lipidomic signature also occurs upon the glial disruption of serine palmitoyltransferase activity (*lace^KD^*) [74], suggesting that *de novo* sphingolipid biosynthesis may be impaired when glia lose Pex5. Interestingly, both glial and neuronal *Pex5^KD^* brains had small but significant increases in species bearing very long chain fatty acids (>C20) (Figure 5H).

In contrast to the sphingolipid defects triggered by glial *Pex5^KD^*, neuronal *Pex5^KD^* caused a phospholipid defect characterized by increases in 18:3(omega-3/ω3) bearing fatty-acyl chains in phosphatidylethanolamines (PE) and phosphatidylcholines (PC) (Figure 5I-J), whereas phosphatidylinositols and phosphatidylserines were not affected (Figure S8D). Intriguingly, PE increased following *Pex5* loss in oligodendrocytes, likely in compensation for decreased ether phospholipids[5]. However, we did not detect major changes to alky-ether phospholipids[75] in neuronal *Pex5^KD^* (Figure S8E), but cannot exclude changes to species not scanned.

Lastly, we also profiled cortex glia *Pex5^KD^* brains by LC-MS/MS (Figure S8F-G), and found modest effects on the global brain lipidome, with weak separation of samples by principal component analysis for TAG, SL, and PL lipids (Figure S8I). In contrast to the pan-glial loss of *Pex5,* cortex glial *Pex5^KD^* did not severely reduce d14/d16 sphingolipid ratios, and instead elevated two CPE species (Figure S8H; CPE d14:1/18:0 and d14:2/18:0). These changes are unlikely to account for the lipid droplet phenotype, which may be driven by neutral lipids not included in the scan (i.e., sterol esters). Taken together, these data show that manipulating peroxisomal import in neurons or glia causes strong and distinct phenotypes on the global brain lipidome.

## Discussion

Here, we uncover circadian regulation of glial peroxisomal dynamics and link clock-controlled organelle biology to brain-wide lipid metabolism and sleep behavior. By systematic profiling of peroxisomal import in the brain, we elucidate that cortex glia have enriched and dynamic peroxisomal import early in the morning. The circadian clock and Pex5 autonomously control circadian import into cortex glia, and cortex glial Pex5 is essential for preventing lipid droplet accumulation and sleep loss. Moreover, glial Pex5 is critical for TAG and sphingolipid homeostasis, whereas neuronal Pex5 was critical for specific glycerophospholipids, highlighting cell-type specific outcomes of peroxisomal import loss in the brain.

### An evolutionarily conserved role for Pex5 in glia

Prior investigations discovered key roles for Pex5 in oligodendrocytes[5] and Schwann cells[6], with severe neurodegeneration and demyelination in *Pex5^KO^*oligodendrocyte, and nonautonomous effects on ion channel distribution downstream of lysosomal dysfunction in Schwann cell *Pex5^KO^*. In contrast, *Pex5^KO^* neurons and astrocytes did not cause overt pathology[17]. By tracking peroxisomal import and conducting *Pex5* knockdown experiments across diverse brain cells, we found that cortex glia are selectively enriched for Pex5-dependent circadian peroxisomal import. Thus, in both flies and humans, glia play a central role in peroxisome-mediated processes, and circadian peroxisomal import should be investigated in mammalian glia which similarly enwrap neuronal soma. Homologues of the cortex glia based on structural morphology include satellite glia cells, protoplasmic astrocytes, and perineuronal oligodendrocytes[76],[77],[78],[79]. Notably, cortex glia extend fine membrane processes tiling dozens of neuronal cell soma, regulate neruonal activity[29],[31], and serve as reservoirs of energy and neutral lipids during development[80],[33] and adulthood[81]. Due to the close association of cortex glia with the neuronal cell soma, we speculate that peroxisomes are vital to neuronal homeostasis through metabolic and cellular detoxifying functions[82],[83],[29]. While we could not detect major lipid changes using LC-MS/MS upon cortex glial *Pex5^KD^*, this may be due to the manipulation targeting ∼2% of the total brain[84]. In contrast, pan-glial *Pex5^KD^* severely depleted TAGs and altered sphingolipid profiles, pointing to important roles for glial peroxisomal import in overall brain lipid metabolism. Future work should determine more precisely how circadian peroxisomal import by cortex glial regulates glial and neuronal metabolism and function.

### Circadian control of peroxisomal import in cortex glia

Circadian clocks regulate daily rhythms in organelle dynamics[52],[85], lipid metabolites[86],[87],[88], and transcriptomes in various tissues[89],[90],[91],[92],[93],[94],[95]. Our work raises the critical question of how pervasively peroxisomal proteins and functions vary across circadian time. While we demonstrate that Pex5 and peroxisomal import peak early in the morning at ZT2 and have a trough at ZT8, future studies should test if all or subsets of peroxisomal proteins are similarly circadian in cortex glia. Surprisingly, despite neurons containing high levels of peroxisomal matrix proteins and PTS1 puncta, we did not detect variations in import across time. Why do cortex glia undergo circadian peroxisomal import? The afternoon cessation of peroxisomal import at ZT8 could suggest a restorative break, as ZT8 aligns with an afternoon siesta thought to adjust for the hottest time of the day[96]. These restorative breaks may be essential for clearing damaged peroxisomes or for timing lipid metabolic loops. However, import does not correlate with activity, arguing against a direct correlation between sleep pressure and peroxisomal import. An alternative hypothesis would be a division of labor in the peroxisome, where at certain times of day specific functions like detoxification or lipid metabolism are favored by enhanced import via Pex5.

While we link the circadian clock to glial peroxisomal import, *period* mutant larval fat bodies have ultradian variation in lipid metabolism and peroxisomal import and biogenesis[97], suggesting both clock and non-clock based mechanisms at play. Intriguingly, a putative deubiquitinase for Pex5, USP2, shows circadian fluctuations in localization to the peroxisome in liver that depend on the core clock[98], and a similar mechanism may regulate the stability and activity of Pex5 in cortex glia.

### Glial peroxisomes in sleep, neurodegeneration and lipid homeostasis

Zellweger spectrum diseases[99],[100] are a group of heterogenous mutations in peroxisomal proteins that cause neurodevelopmental and neurodegenerative disorders.[101] As cortex glial Pex5 is essential for sleep and subject to circadian regulation, it may be fruitful to investigate for circadian modulation of Zellweger disorder phenotypes, including sleep. Our behavioral findings suggest a model wherein cortex glial communicate with closely associated neuronal soma to regulate behavioral processes such as sleep [102],[103]. Sleep deprivation induces a cytokine storm[104], and *Pex5* perturbation also induces pro-inflammatory cytokine *upd3* in glial cells[4]. In flies, altered lipid droplet metabolism in cortex and ensheathing glia downstream of neuronal oxidative stress can cause sleep defects[71]. Moreover, a recent preprint reports that sleep itself can control peroxisomal abundance and downstream oxidative damage in neurons [105]. Given that *Pex5* knockdown in cortex glia caused sleep loss and increased lipid droplets, we speculate that neurons in aged *Pex5^KD^* flies could accumulate reactive oxygen species and upregulate cytokines through metabolic coupling between neurons and glia.

Knockdown of Pex5 caused cell type-specific alterations to the young adult brain lipidomes, arguing that peroxisomes serve different metabolic functions in neurons (phospholipid homeostasis) versus glia (sphingolipid and TAG metabolism). The broad loss of TAG upon glial knockout may occur from increased lipolysis of lipid droplets[106] and/or decreased neutral lipid uptake and storage, which could be direct or secondary effects of Pex5 depletion. Enhanced lipolysis may reflect a shifting energy balance of glial cells with compromised peroxisomal import, and glial mitochondrial beta-oxidation is a key determinant of both fly viability and TAG and lipid droplet abundances[107]. Prior work on global *pex2, pex3,* and *pex16* mutants also observed changes in sphingolipid homeostasis[108],[109]. The surprising phenotypic overlap of decreased d14, and increased d16, sphingolipids in glial *Pex5* knockdown (Figure 5D-H) and glial knockdown of *lace*, the rate limiting enzyme in sphingolipid biosynthesis, suggests that *Pex5* deficient glia are defective in *de novo* sphingolipid biosynthesis. Whether this effect is primary (defects in fatty acid substrates) or secondary (compromised glial health) remains to be determined. Lastly, we were surprised to observe higher n-3 phosphatidylethanolamines in neuronal *Pex5* knockdown brains, given that Zellweger patients often lose n-3 glycerophospholipids[110]. While our data point to important roles for neuronal peroxisomes in phospholipid metabolism, future work is required to decipher the cause, and consequences, of these lipid alterations.

In conclusion, by studying Pex5 in the adult *Drosophila* brain and the dynamics of peroxisomal import, we identify extensive cell-type specific variability, with elevated import in neurons and cortex glia. In cortex glia, import and Pex5 levels are under strict circadian control. Our identification of sleep disruption upon Pex5 loss in cortex glia opens the avenue to a deeper understanding of peroxisome function in the brain, and the neurological consequences of peroxisomal disorders.

---

**Figure S1, related to Figure 1.**
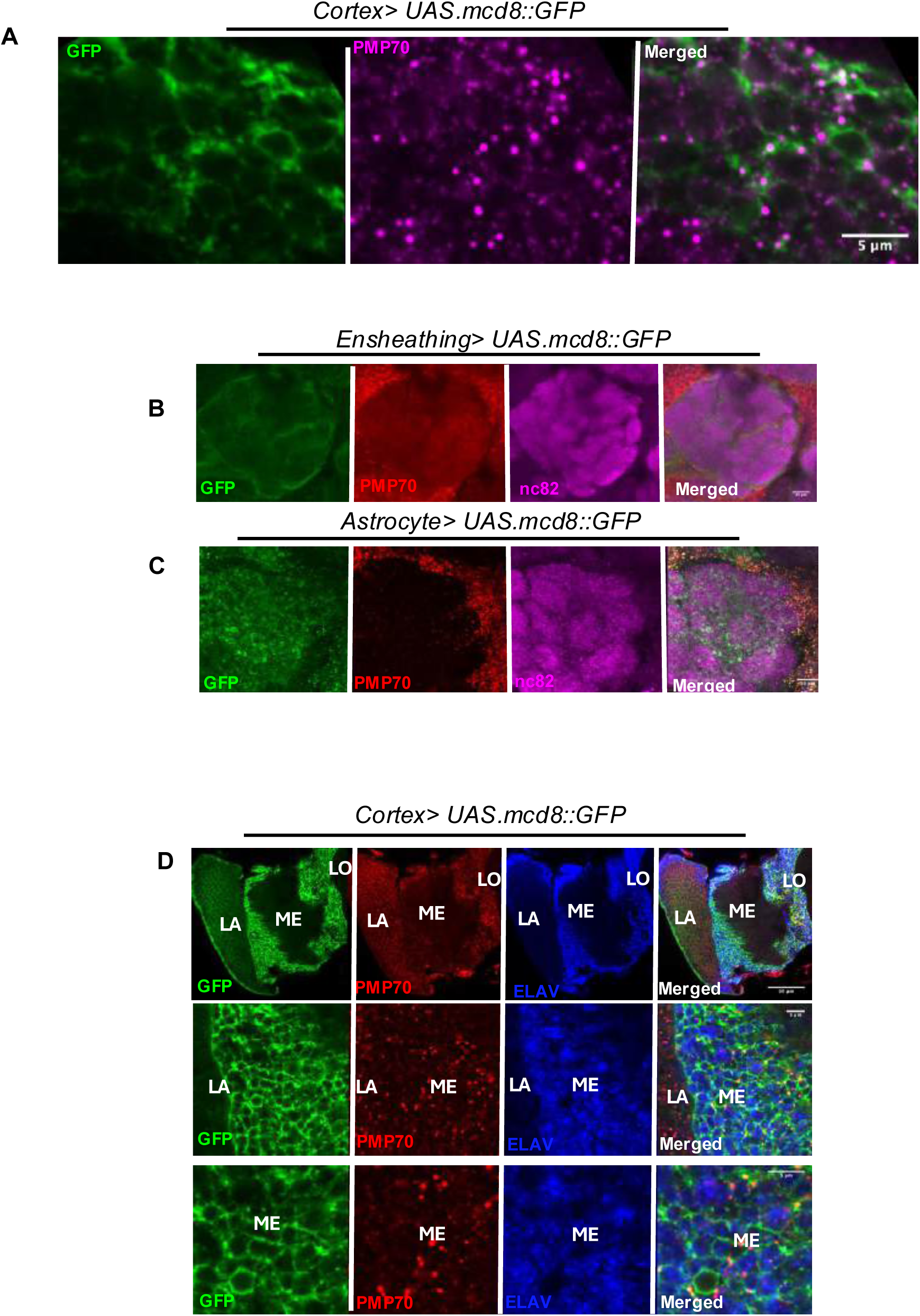
Representative images comparing central brain glial classes, with colocalization between glial driven GFP(green) counterstained with PMP70(magenta). PMP70 peroxisomes are abundant inside cortex glial regions as compared to astrocyte and ensheathing glia. nC82(magenta) marks the pre-synaptic active zones representing the neuropils of the adult *Drosophila* brain. **(A)** Adult female *Drosophila* brains (7 days old) Cortex glia (*GMR54H02*-GAL4> UAS-*mCD8::GFP*) tracking colocalization between GFP(green) counterstained with PMP70(magenta). **(B)** Adult female *Drosophila* brains (7 days old) Ensheathing glia (*GMR56F03-GAL4*> UAS-*mCD8::GFP*) tracking colocalization between GFP(green) counterstained with PMP70(red) and nc82 (magenta). **(C)** Adult female *Drosophila* brains (7 days old) astrocytic glia (*alrm-GAL4> UAS-mCD8::GFP*) tracking colocalization between GFP(green) counterstained with PMP70(red) and nc82 (magenta). **D)** Adult female *Drosophila* brains (7 days old) Cortex glia (*GMR54H02*-GAL4> UAS-*mCD8::GFP*) representative image showing the optic lobe regions (LA-lamina; ME-medulla; LO-lobula) showing PMP70 (red) positive puncta in GFP(green) rich regions of cortex.

**Figure S2, related to Figure 2.**
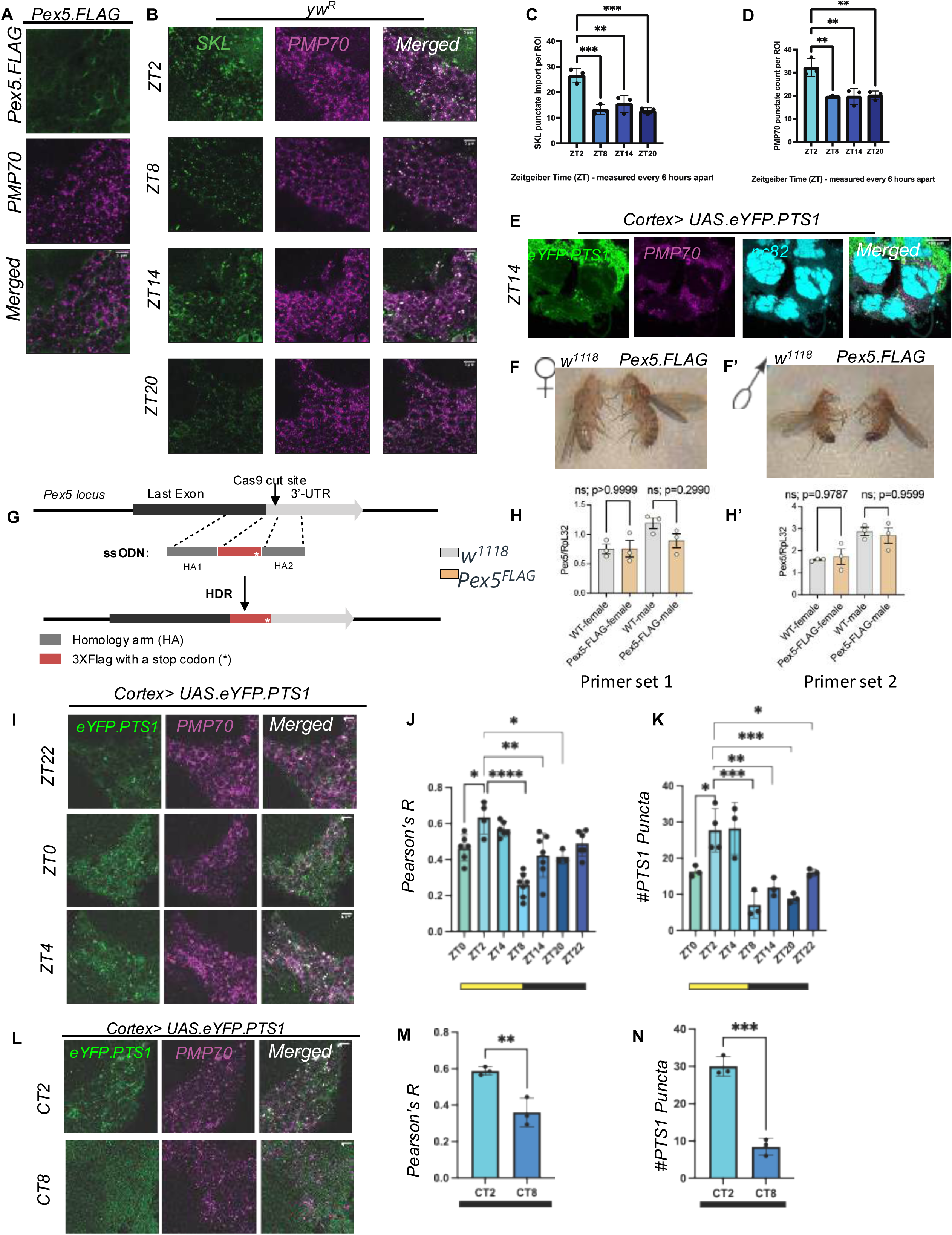
**(A)** Adult female *Drosophila* brains (7 days old) tracking Pex5.FLAG knock-in flies co-stained with anti-FLAG(green) and anti-PMP70(magenta). **(B)** Adult female *Drosophila* brains (7 days old) *yw^R^* wildtype flies dissected across ZT2, ZT8, ZT14 and ZT20 and stained with anti-SKL(green), anti-PMP70(magenta) to show circadian oscillations of peroxisomal import peak at ZT2 and trough at ZT8. **(C)** Quantification in *yw^R^* wildtype flies of *SKL* puncta at ZT2, ZT8, ZT14, ZT20. *n* ≥*3* brains per genotype, dots represent biological replicates averaged over ≥*3* ROIs. ROIs were taken from zooms into cortex below the antennal lobes and above the suboesophageal ganglion. **(D)** Quantification in *yw^R^* wildtype flies of PMP70 puncta at ZT2, ZT8, ZT14, ZT20; *n* ≥*3* brains per genotype, dots represent biological replicates averaged over ≥*3* ROIs. **(E)** Adult female *Drosophila* brains (7 days old) tracking peroxisomal import oscillation at ZT14 observed in glia with *UAS-eYFP.PTS1* using *GMR54H02-GAL4*. eYFP-tagged *PTS1*-protein import (green) is counterstained with PMP70 (magenta), an abundant peroxisomal membrane protein to detect peroxisomes along with anti-nc82. Scale bar = 50μm. **(F)** Adult female *Drosophila* comparing *w^1118^* versus *Pex5.FLAG*. **(F’)** Adult male *Drosophila* comparing *w^1118^* versus *Pex5.FLAG*. **(G)** CRISPR/Cas9-based knock-in strategy to tag *Pex5* with a 3XFLAG epitope. **(H)** Relative transcript expression of Pex5 (Primer set 1) in w1118 versus Pex5.FLAG measured by RT-qPCR (full flies, both adult female and male; 7 days old) at ZT2. **(H’)** Relative transcript expression of Pex5 (Primer set 2) in w1118 versus Pex5.FLAG measured by RT-qPCR (full flies, both adult female and male; 7 days old) at ZT2. **(I)** Adult female *Drosophila* brains (7 days old) *Cortex>UAS-eYFP.PTS1* flies dissected across ZT22, ZT0 and ZT4 stained with anti-GFP(green), anti-PMP70(magenta). **(J)** Quantification of eYFP-PTS1 and PMP70 Pearson colocalization. *n* ≥*3* brains per genotype, data points represent biological replicates. **(K)** Quantification of the number of PTS1 punctate. *n* ≥*3* brains per genotype, dots represent biological replicates averaged over ≥*3* ROIs. **(L)** Adult female *Drosophila* brains (7 days old) *Cortex>UAS-eYFP.PTS1* flies dissected across CT2 and CT8 (CT = circadian time) in 12hr/2hr dark/dark cycle after initial light/dark entrainment stained with anti-GFP(green), anti-PMP70(magenta). **(M)** Quantification of eYFP-PTS1 and PMP70 Pearson colocalization. *n* ≥*3* brains per genotype, data points represent biological replicates. **(N)** Quantification of the number of PTS1 punctate. *n* ≥*3* brains per genotype, dots represent biological replicates averaged over ≥*3* ROIs. ROIs were taken from zooms into cortex below the antennal lobes and above the suboesophageal ganglion. ns= non-significant, * p < 0.05, ** p < 0.01, *** p < 0.001, **** p < 0.0001 by Tukey’s multiple comparisons test.

**Figure S3, related to Figure 2.**
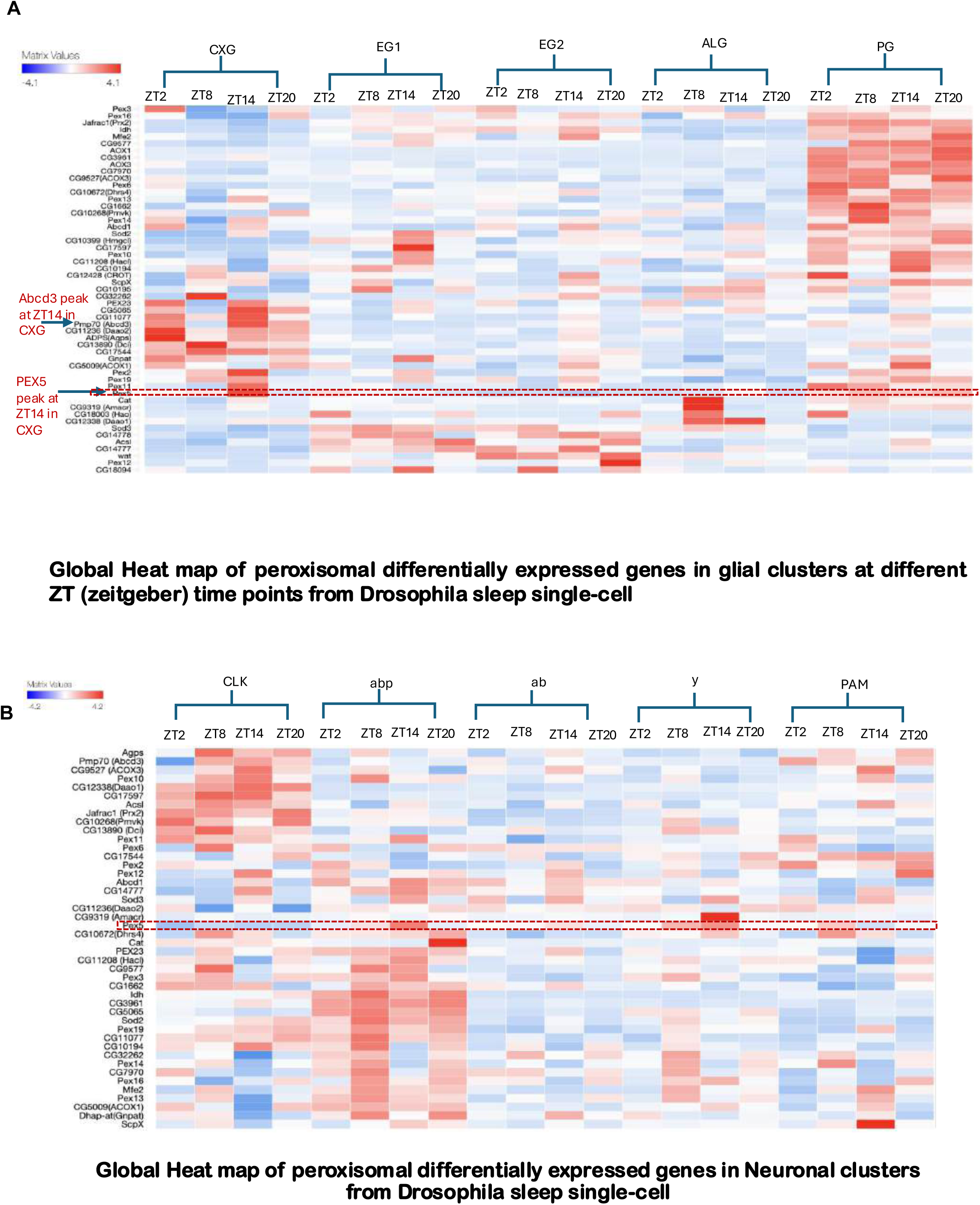
**(A)** Global heatmap of peroxisomal gene expression extracted from fly-sleep single across all glial clusters to show *Pex5* expression is highest in CXG (cortex-glia) cluster. **(B)** Global heatmap of peroxisomal gene expression extracted from fly-sleep single across neuronal clusters to show low *Pex5* expression as opposed to CXG glial cluster.

**Figure S4, related to Figure 2.**
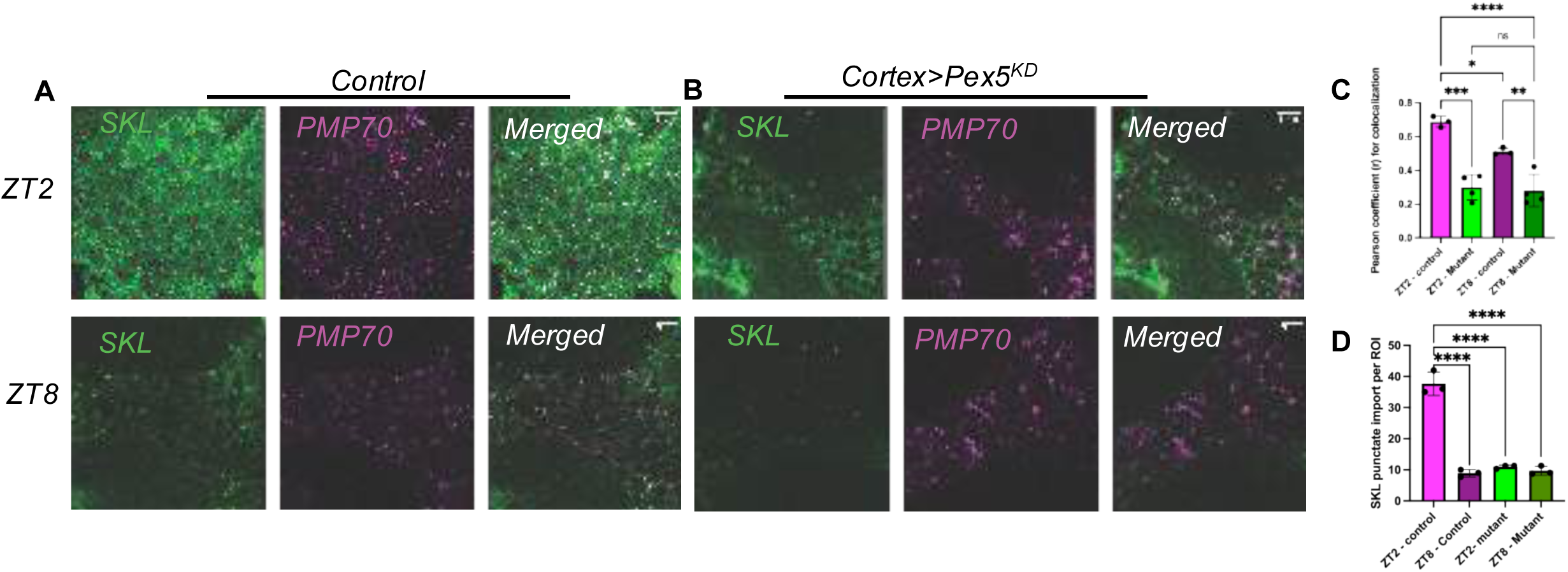
**(A)** Adult female *Drosophila* brains (7 days old) Control (*w^1118^ /UAS-Pex5* RNAi) flies dissected across ZT2 and ZT8 stained with anti-SKL(green), anti-PMP70(magenta) to show circadian oscillations of peroxisomal import peak at ZT2 and trough at ZT8. **(B)** Adult female *Drosophila* brains (7 days old) mutants (*Cortex > UAS-Pex5 RNAi;* (BDSC# 55322) flies dissected across ZT2 and ZT8 stained with anti-SKL(green), anti-PMP70(magenta). **(C)** Quantification of SKL and PMP70 Pearson colocalization. *n* ≥*3* brains per genotype, data points represent biological replicates. ROIs were taken from zooms into cortex below the antennal lobes and above the suboesophageal ganglion. **(D)** Quantification of the number of SKL punctate. *n* ≥*3* brains per genotype, dots represent biological replicates averaged over ≥*3* ROIs. * p < 0.05, ** p < 0.01, *** p < 0.001, **** p < 0.0001 by Tukey’s multiple comparisons test.

**Figure S5, related to Figure 3.**
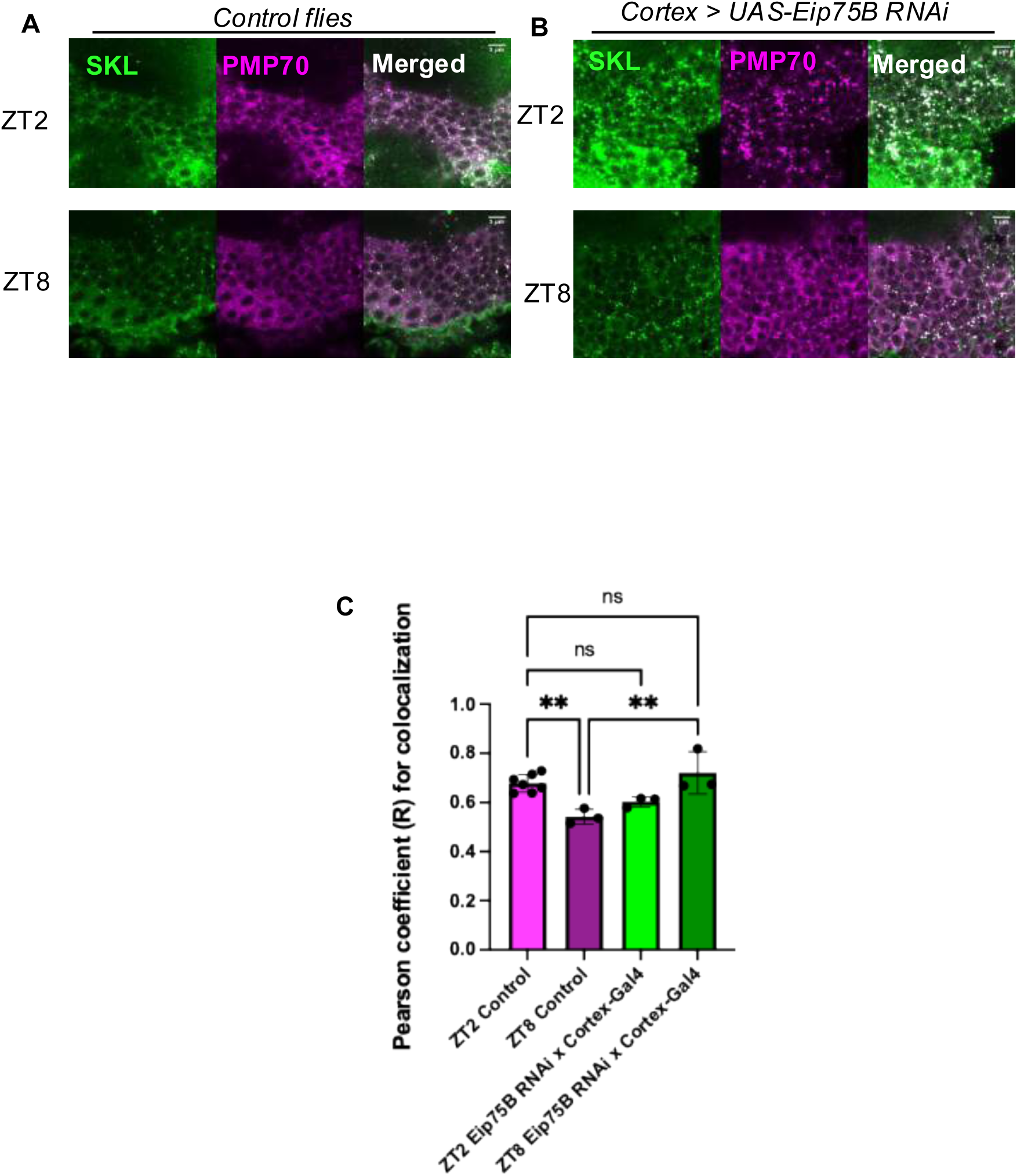
**(A)** Adult female *Drosophila* brains (7 days old) dissected at ZT2 & ZT8 for control flies (*w^1118^/UAS-Eip75B RNAi*) stained with anti-SKL(green) and anti-PMP70(magenta). **(B)** Adult female *Drosophila* brains (7 days old) dissected at ZT2 & ZT8 for *GMR54H02-Gal4>UAS-Eip75B RNAi* knockdown flies. **(C)** Quantification Pearson R coefficient colocalization calculated (see methods) for *GMR54H02-Gal4* cross with *UAS-Eip75B* RNAi knockdown flies compared with control flies at ZT2, ZT8; *n* ≥*3* brains per genotype, dots represent biological replicates averaged over ≥*3* ROIs. ROIs were taken from zooms into cortex below the antennal lobes and above the suboesophageal ganglion, * p < 0.05, ** p < 0.01, *** p < 0.001, **** p < 0.0001 by Tukey’s multiple comparisons test.

**Figure S6, related to Figure 3.**
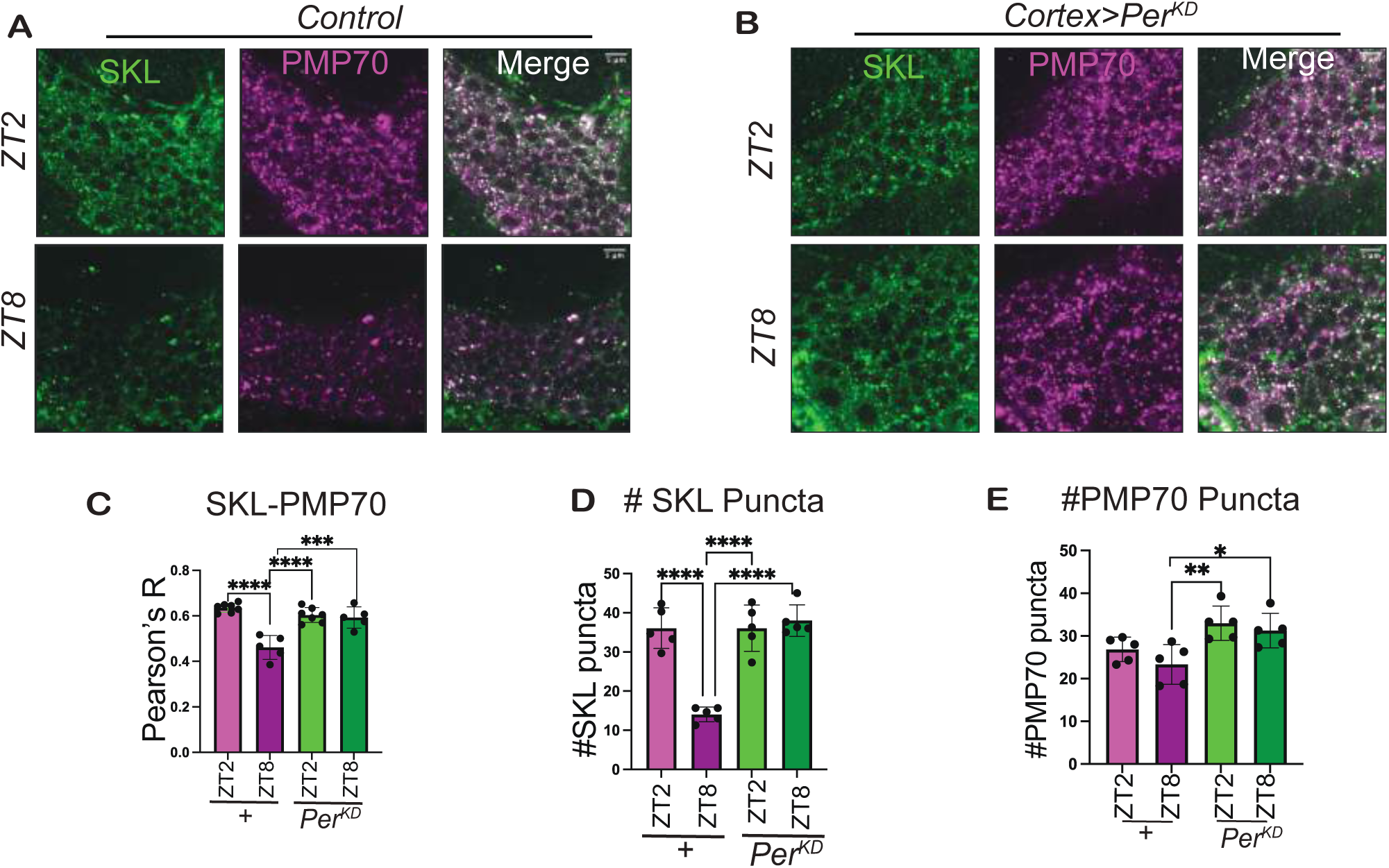
Per RNAi autonomous control in cortex glia. **(A)** Adult female *Drosophila* brains (7 days old) tracking peroxisomal import oscillation across ZT2, ZT8 in Control *>w1118/GMR54H02-Gal4)* with SKL (green) counterstained with PMP70 (magenta). **(B)** Adult female *Drosophila* brains (7 days old) tracking peroxisomal import oscillation across ZT2, ZT8 in *Cortex-Glia> UAS-Per RNAi* (*GMR54H02-Gal4>UAS-Per RNAi)* with SKL (green) counterstained with PMP70 (magenta). **(C)** Quantification of SKL and PMP70 Pearson colocalization. *n* ≥*3* brains per genotype, data points represent biological replicates averaged over ≥*3* ROIs. ROIs were taken from zooms into cortex below the antennal lobes and above the suboesophageal ganglion, **(D)** Quantification of the number of SKL punctate. *n* ≥*3* brains per genotype, dots represent biological replicates averaged over ≥*3* ROIs. **(E)** Quantification of the number of PMP70 punctate. *n* ≥*3* brains per genotype, dots represent biological replicates averaged over ≥*3* ROIs. * p < 0.05, ** p < 0.01, *** p < 0.001, **** p < 0.0001 by Tukey’s multiple comparisons test.

**Figure S7, related to Figure4.**
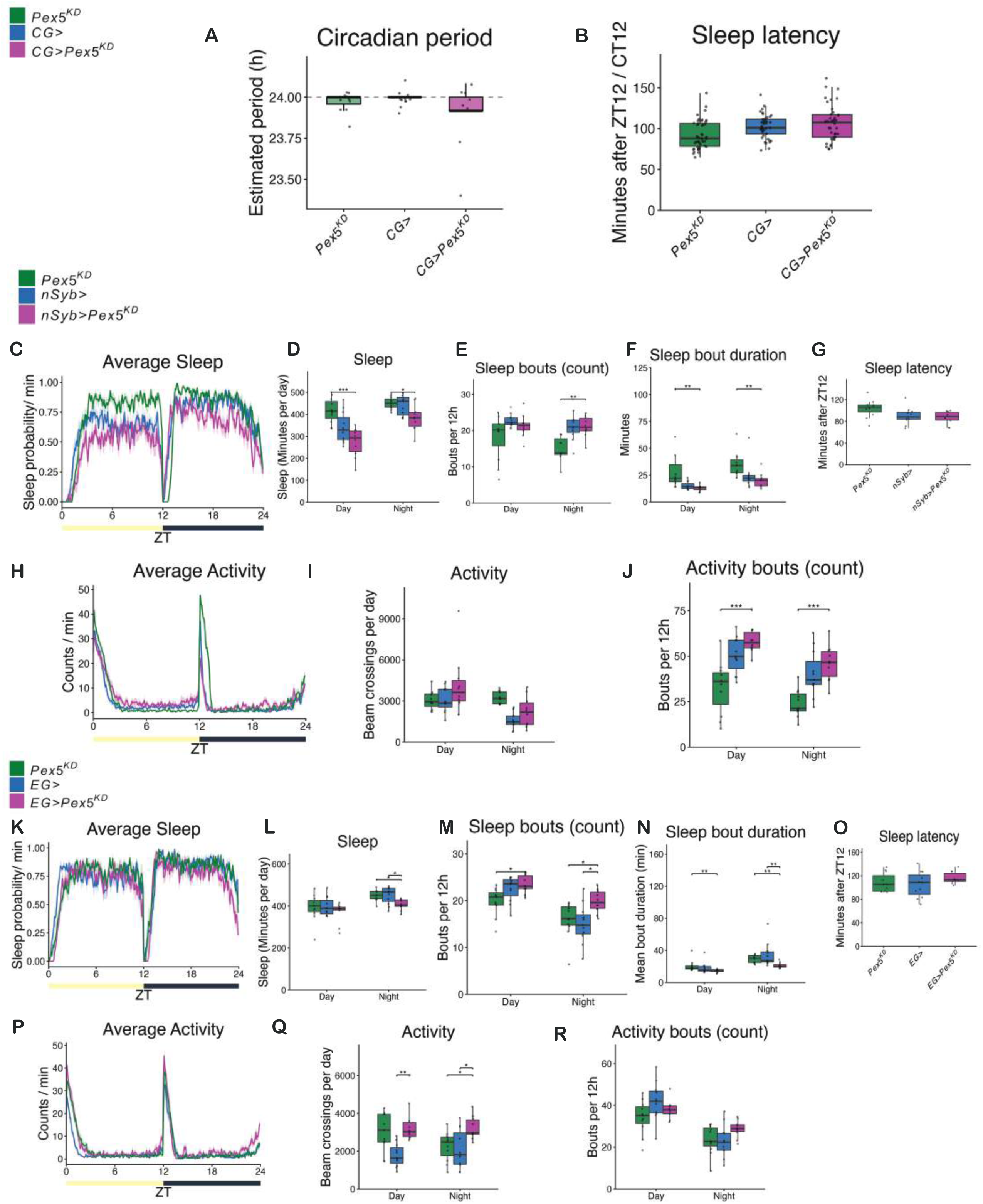
Loss of *Pex5* sleep and activity analyzed for neuronal and ensheathing glia. **(A)** Circadian period length measured in 12hr/12hr, dark/dark cycle on an average for 5-6 days continuously for 5-day to 10-day old males post-eclosion after initial entrainment in 12hr/12hr, light/dark for cortex glia driver *GMR77A03-GAL4* crossed with *UAS-Pex5 RNAi KD* (magenta) compared to parental controls *UAS-Pex5-RNAi (*green) *& GMR77A03-GAL4 (blue)*. **(B)** Sleep latency was measured after lights off in 12hr/12hr, light/dark cycle on an average for 4-5 days continuously for 5-day to 10-day old males post-eclosion for cortex glia driver *GMR77A03-GAL4* crossed with *UAS-Pex5 RNAi KD* (magenta) compared to parental controls *UAS-Pex5-RNAi (*green) *& GMR77A03-GAL4 (blue)*. **(C)** Average Sleep (sleep probability/min) measured in 12hr/12hr, light/dark cycle on an average for 4-5 days continuously for 5-day to 10-day old males post-eclosion. Average sleep time-plot showing sleep probability/min (bouts of inactivity > 5 mins) for the neuronal driver *nSyb-GAL4* crossed with *UAS-Pex5 RNAi KD* (magenta) compared to parental controls *UAS-Pex5-RNAi (*green) *& nSyb-GAL4 (blue)*. **(D)** Total sleep(minutes per day) quantified for all three genotypes from **(C).** Individual black dots represent different biological replicates. **(E)** Sleep bouts per 12hour quantified for all three genotypes from **(C).** Individual black dots represent different biological replicates. **(F)** Sleep bout duration per minute quantified for all three genotypes from **(C).** Individual black dots represent different biological replicates. **(G)** Sleep latency quantified for all three genotypes from **(C).** Individual black dots represent different biological replicates. **(H)** Average activity measured in 12hr/12hr, light/dark cycle on an average for 4-5 days continuously for 5-day to 10-day old males post-eclosion. Average activity time-plot showing counts/min (bouts of inactivity > 5 mins) for the neuronal driver *nSyb-GAL4* crossed with *UAS-Pex5 RNAi KD* (magenta) compared to parental controls *UAS- Pex5-RNAi (*green) *& nSyb-GAL4 (blue)*. **(I)** Total activity (beam crossings per day) quantified for all three genotypes from **(H).** Individual black dots represent different biological replicates. **(J)** Activity bouts (per 12 hour) quantified for all three genotypes from **(H).** Individual black dots represent different biological replicates. **(K)** Average Sleep (sleep probability/min) measured in 12hr/12hr, dark/dark cycle on an average for 5-6 days continuously for 5-day to 10-day old males post-eclosion after initial entrainment on light/dark cycle. Average sleep time-plot showing sleep probability/min (bouts of inactivity > 5 mins) for the ensheathing glia driver *GMR56F03-GAL4* crossed with *UAS-Pex5 RNAi KD* (magenta) compared to parental controls *UAS-Pex5- RNAi (*green) *& GMR56F03-GAL4 (blue)*. **(L)** Total sleep(minutes per day) quantified for all three genotypes from **(K).** Individual black dots represent different biological replicates. **(M)** Sleep bouts per 12hour quantified for all three genotypes from **(K).** Individual black dots represent different biological replicates. **(N)** Sleep bout duration per minute quantified for all three genotypes from **(K).** Individual black dots represent different biological replicates. **(O)** Sleep latency quantified for all three genotypes from **(K).** Individual black dots represent different biological replicates. **(P)** Average activity measured in 12hr/12hr, dark/dark cycle on an average for 5-6 days continuously for 5-day to 10-day old males post-eclosion after initial entrainment on light/dark cycle. Average activity time-plot showing counts/min (bouts of inactivity > 5 mins) for the ensheathing glia driver *GMR56F03-GAL4* crossed with *UAS-Pex5 RNAi KD* (magenta) compared to parental controls *UAS-Pex5-RNAi (*green) *GMR56F03-GAL4 (blue)*. **(Q)** Total activity (beam crossings per day) quantified for all three genotypes from **(P).** Individual black dots represent different biological replicates. **(R)** Activity bouts (per 12 hour) quantified for all three genotypes from **(P).** Individual black dots represent different biological replicates. * p < 0.05, ** p < 0.01, *** p < 0.001, **** p < 0.0001 by ANOVA, Tukey’s multiple comparisons or Welch’s ANOVA depending on equality of data variance. For nonparametric sleep bouts, Mann-Whitney U test was used. n ≥ 10 flies for light/dark experiment.

**Figure S8, related to Figure5.**
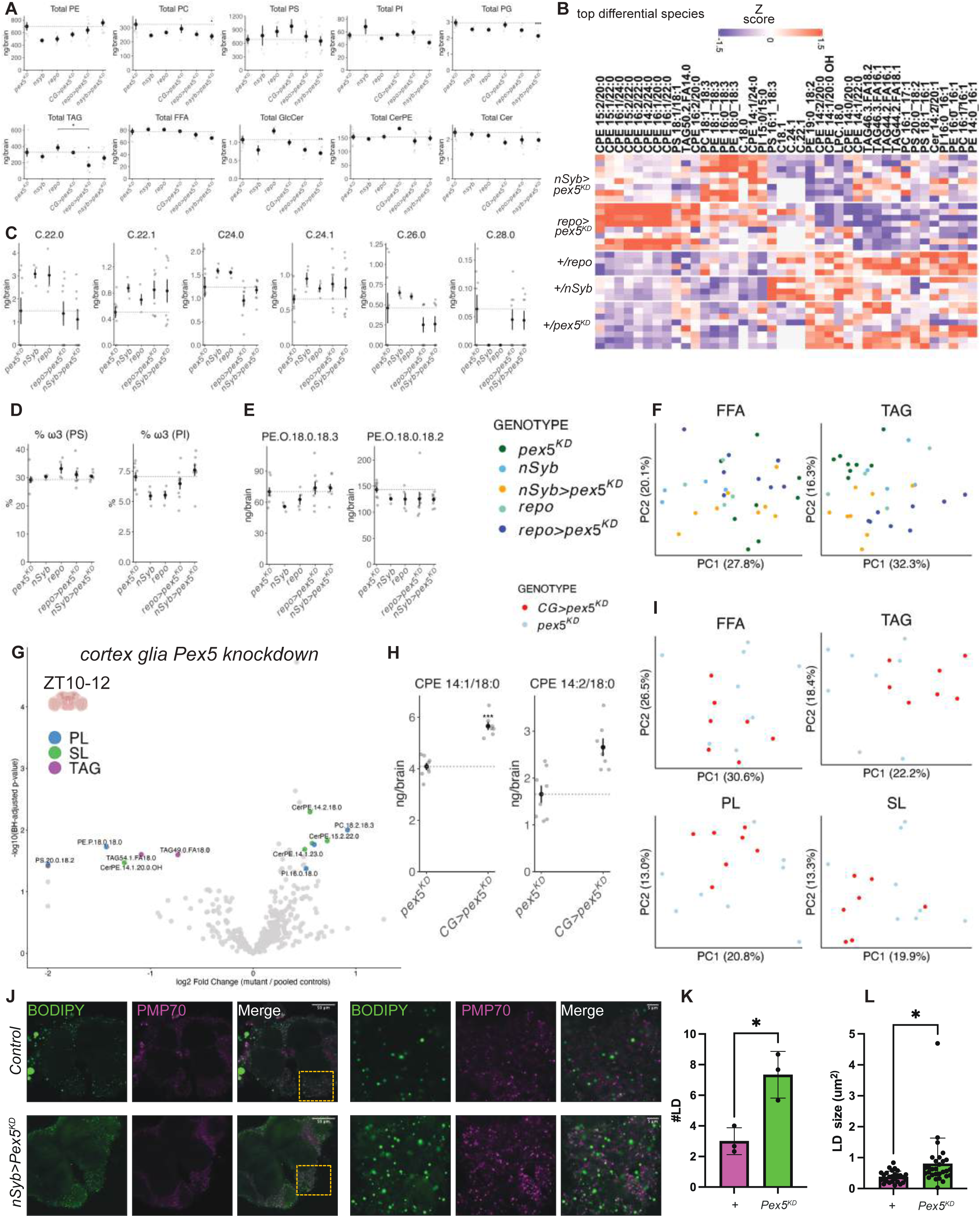
Pex5 is essential for lipid homeostasis in the adult brain. **(A)** Total ng/brain for lipid families analyzed by LC-MS/MS of day 7 females brains at ZT10-12; see Table S1 for full genotypic details. *n* ≥*4* tubes of 20 brains for each genotype. **(B)** Heatmap of top 30 species changed between controls and mutants, z-scored by relative % abundance. **(C)** Free fatty acid analysis of longer chain fatty acid species across genotype, including very long chain fatty acids (VLCFA) (ng/brain) **(D)** % of ω3 fatty acids in phosphatidylserine (PS) and phosphatidylinositol (PI). **(E)** Major alky-ether phospholipids shown in ng/brain across genotypes. **(F)** Principal component analysis of TAGs and free fatty acids (FFA) colored by genotype. **(G)** Volcano plot showing p-value versus log2 fold change for cortex glia *pex5* knockdown versus parental controls. Phospholipids are colored green, sphingolipids colored green, and TAG colored purple. **(H)** Differentially abundant ceramide phosphoethanolamine (CPE) species in *CG > pex5^KD^* brains. **(I)** principal component analysis of controls versus *CG > pex5^KD^* brains (red) versus control (grey) for sphingolipids (SL), phospholipids (PL), TAG, and free fatty acids (FFA). **(J)** Adult female *Drosophila* brains (7 days old) tracking neutral lipid droplet (BODIPY 493/503, green) counterstained with PMP70 (magenta). Representative images of UAS-control (*UAS-Pex5 RNAi > w^1118^*) versus neuronal specific knockdown of Pex5 using *nSyb-Gal4* > *UAS-Pex5 RNAi*. **(K)** Quantification of different lipid droplet size(um^2^) from J. *n* ≥*3* brains per genotype, dots represent individual lipid droplets per biological replicates averaged over ≥*3* ROIs. ROIs were taken from zooms into cortex below the antennal lobes and above the suboesophageal ganglion, and quantified from 200×200 pixel windows inside these zooms. **(L)** Quantification of lipid droplet count from J. *n* ≥*3* brains per genotype, dots represent number of lipid droplets per biological replicates averaged over ≥*3* ROIs. * p < 0.05, ** p < 0.01, *** p < 0.001, **** p < 0.0001 by Tukey’s multiple comparisons test.

## RESOURCE AVAILABILITY

### Lead contact

Correspondence should be directed to Anurag Das

### Materials availability

All data and code reported in this paper will be shared by the lead contact (AD) upon request.

## Acknowledgements

We thank Bai lab members for feedback during lab meetings. We thank Bloomington Drosophila Stock Center (NIH P40OD018537) for fly stocks. We thank FlyBase (NHGRI U41HG000739) for *Drosophila* gene annotation. We thank Kyu-Sun Lee for fly Pmp70 antibodies. We thank Abhilash Goswami for providing suggestions for the extraction of peroxisomal genes from publicly available single cell data. We thank Elizabeth McNeill, Amita Sehgal, and Marc Tatar for sharing fly stocks. We thank Alicia Taylor from the Elizabeth McNeill Lab for providing DAM5H-Trikinetics Drosophila Sleep Activity Monitoring system for our sleep studies. Graphical model figures were created with BioRender.com. This work was supported by NSF CAREER 2046984, Hevolution HF-GRO-23-1199062-14, and NIH R01AG075156 to HB. JPV is supported by the Sandler PBBR grant and the Simons Foundation.

## Author contributions

**Conceptualization:** Anurag Das, Hua Bai

**Data Curation:** Anurag Das, John Philip Vaughen

**Formal Analysis:** Anurag Das, John Philip Vaughen, Hua Bai

**Funding Acquisition:** John Philip Vaughen, Vera Mazurak, Hua Bai

**Investigation:** Anurag Das, John Philip Vaughen, Irma Magaly Rivas-Serna, Marlene Dorneich-Hayes, Lakpa Sherpa, Hia Kalita

**Methodology:** Anurag Das, John Philip Vaughen, Irma Magaly Rivas-Serna, Ankur Kumar, Ruiqi Liu, Kerui Huang, Hua Bai

**Project Administration:** Anurag Das, John Philip Vaughen, Hua Bai

**Resources:** Anurag Das, John Philip Vaughen, Vera Mazurak, Hua Bai

**Supervision:** John Philip Vaughen, Hua Bai

**Validation:** Anurag Das, Hua Bai, John Philip Vaughen, Lakpa Sherpa, Hia Kalita

**Visualization:** Anurag Das, John Philip Vaughen, Hua Bai

**Writing - Original Draft Preparation:** Anurag Das

**Writing - Review & Editing:** Anurag Das, John Philip Vaughen

**Revisions:** Anurag Das, John Philip Vaughen

## Resources table

| REAGENT or RESOURCE | SOURCE | IDENTIFIER |
| --- | --- | --- |
| Antibodies |  |  |
| Fly-PMP70 | a gift from Kyu-Sun Lee | N/A |
| Anti-ELAV | DSHB | Rat-Elav-7E8A10 anti-elav |
| Anti-nc82 | DSHB | nc82 |
| Monoclonal Anti-FLAG | Sigma-Aldrich | F1804 |
| Anti-SKL | a gift from Richard Rachubinski | N/A |
| Anti-GFP mAb | Cell Signaling Technology | 2956 |
| Experimental models: Organisms/strains |  |  |
| *D. melanogaster*: *w*<sup>1118</sup> | BDSC | Cat# 5905; RRID: BDSC_5905 |
| *D. melanogaster*: *ywR* | A gift from Marc Tatar | N/A |
| *D. melanogaster*: *CantonS* | A gift from Elizabeth McNeill | N/A |
| *D. melanogaster*: *per*[01] | BDSC | Cat# 80928; RRID: BDSC_80928 |
| *D. melanogaster*: *w*[1118]; *P*{*y*[+*t*7.7]<br>*w*[+*mC*]=*GMR77A03-GAL4*}*attP2* | BDSC | Cat# 39944; RRID: BDSC_39944 |
| *D. melanogaster*: *w*[1118]; *P*{*y*[+*t*7.7]<br>*w*[+*mC*]=*GMR54H02-GAL4*}*attP2* | BDSC | Cat# 45784; RRID: BDSC_45784 |
| *D. melanogaster*: *y*[1] *v*[1]; *P*{*y*[+*t*7.7]<br>*v*[+*t*1.8]=*TRiP.HMJ21920*}*attP40* | BDSC | Cat# 58064; RRID: BDSC_58064 |
| *D. melanogaster*: *w*[1118]; *P*{*y*[+*t*7.7]<br>*w*[+*mC*]=*GMR57C10-GAL4*}*attP2* | BDSC | Cat# 39171; RRID: BDSC_39171 |
| *D. melanogaster*: *y*[1] *sc*[*] *v*[1] *sev*[21];<br>*P*{*y*[+*t*7.7] *v*[+*t*1.8]=*TRiP.HMS01530*}*attP40* | BDSC | Cat# 35780; RRID: BDSC_35780 |
| *D. melanogaster*: *w*[1118]; *P*{*y*[+*t*7.7]<br>*w*[+*mC*]=*GMR56F03-GAL4*}*attP2* | BDSC | Cat# 39157; RRID: BDSC_39157 |
| *D. melanogaster*: *w*[1118]; *P*{*y*[+*t*7.7]<br>*w*[+*mC*]=*GMR83E10-GAL4*}*attP2* | BDSC | Cat# 40362; RRID: BDSC_40362 |
| *D. melanogaster*: *Pex5*[*FLAG-a1*]/*FM7a* | This paper | N/A |
| *D. melanogaster*: *w* [1118];<br>*P* { *w* [+ *m* *]= *GAL4* } *repo* /TM3, *Sb* [1] | BDSC | Cat# 7415; RRID: BDSC_7415 |
| *D. melanogaster*: *w*[*]; *P*{*w*[+*mC*]=*UAS-*<br>*eYFP.PTS1*}6 | BDSC | Cat# 64248; RRID: BDSC_64248 |
| *D. melanogaster*: *y*[1] *w*[*]; *betaTub60D*[*Pin-*<br>*Yt*]/*CyO*; *P*{*w*[+*mC*]=*UAS-mCD8::GFP.L*}*LL6* | BDSC | Cat# 5130; RRID: BDSC_5130 |
| *D. melanogaster*: *w* [*]; *wg* [*Sp-1*]/ *CyO*;<br>*P* { *w* [+ *m* *]= *alrm-GAL4.D* }2 | BDSC | Cat# 67032; RRID: BDSC_67032 |
| *D. melanogaster*: *y*[1] *v*[1];<br>*P*{*y*[+*t*7.7]=*CaryP*}*Msp300*[*attP40*] | BDSC | Cat# 36304; RRID: BDSC_36304 |
| *D. melanogaster*: *repo-GAL4* = +; *UAS-Dcr-2*<br>(BDSC 24650)/+; *repo-GAL4* (BDSC 7415)/+ | This paper | N/A |
| *D. melanogaster*: *nSyb-GAL4* = +; *UAS-Dcr-*<br>*2*/+; *nSyb-GAL4* (BDSC 39171)/+ | This paper | N/A |
| *D. melanogaster*: *repo* > *pex5*<sup>KD</sup> = +; *UAS-*<br>*Dcr-2/pex5-trip*; *repo-GAL4*/+ | This paper | N/A |
| *D. melanogaster*: CG > *pex5*<sup>KD</sup> = +; *pex5-trip*/+; GMR54H02-GAL4(BDSC 45784)/+ | This paper | N/A |
| *D. melanogaster*: *nSyb* > *pex5*<sup>KD</sup> = +; UAS-*Dcr-2/pex5-trip*; *nSyb*-GAL4/+ ; | This paper | N/A |
| *D. melanogaster*: *y[1] sc[\*] v[1] sev[21]*; *P{y[+t7.7]v[+t1.8]=TRiP.HMS02045}attP2/TM 3, Sb[1]* | BDSC | Cat# 40878; RRID: BDSC_40878 |
| *D. melanogaster*: *w[\*]*; *P{w[+mC]=UAS-Clk.Delta}1* | BDSC | Cat# 36318; RRID: BDSC_36318 |
| *D. melanogaster*: *w[1118]*; *P{w[+mC]=UAS-Dcr-2.D}2* | BDSC | Cat# 24650; RRID: BDSC_24650 |
| *D. melanogaster*: *w[1118]*; *PBac{y[+mDint2] GFP[E.3xP3]=vas-Cas9}VK00027* | BDSC | Cat# 51324; RRID: BDSC_51324 |
| *D. melanogaster*: *FM7i*, *P{w[+mC]=ActGFP}JMR3/C(1)DX, y[1] f[1]* | BDSC | Cat# 4559; RRID: BDSC_4559 |
| *D. melanogaster*: *y[1] v[1]*; *P{y[+t7.7] v[+t1.8]=TRiP.HMC04009}attP40* | BDSC | Cat# 55322; RRID: BDSC_55322 |
| Other |  |  |
| BODIPY 493/503 Thermo Fisher Scientific | Thermo | Cat# D3922 |
| Software and algorithms |  |  |
| ImageJ | FIJI | [imagej.net (USA)](https://imagej.net) |
| Graphpad Prism 10 | Graph Pad Software | Subscription ID :2814928 |
| ADOBE Illustrator 2020 | ADOBE | N/A |
| Clock Lab Analysis 2 | Actimetrics | N/A |
| R Studio 2023.12.1 (4.3.1) | R | RRID: SCR_000432 |

## EXPERIMENTAL MODEL AND METHOD DETAILS

### Fly Husbandry and Stocks

*Drosophila melanogaster* were maintained at 25°C, 60% relative humidity and 12-hour light/dark cycle unless specified otherwise. Adults were reared on an agar-based diet consisting of 0.8% cornmeal, 10% sugar, and 2.5% yeast. Fly stocks used in the present study are mentioned in the above resources table. Female flies (7 days old) were used in lipidomics and confocal experiments, while male flies were used in sleep studies from 6 to 10 days old. Flies and crosses were maintained in plastic vials and bottles in a thermoregulated chamber at 25 degree Celsius following a 12:12 h light-dark cycle with 60% humidity. All experiments were performed 2-3 times independently.

### Pex5.FLAG knock-in

The *Pex5.FLAG* knock-in fly was generated using CRISPR/Cas9-mediated homology-directed repair in *Drosophila* (*Figure S2G*). Briefly, the gRNA targeting the 3’-UTR of the *Pex5* locus was cloned into a *pU6-2-chiRNA* vector. The ssODN with 3XFlag sequences flanking with two homology arms was synthesized by Integrated DNA Technologies Inc. The gRNA vector and ssODN were injected into *w1118* flies expressing *vas-Cas9* (BL51324). Injected G0 flies were crossed to an attached-X stock (BL4559) to generate F1 progeny. F1 males carrying the FLAG insertion were identified via PCR and Sanger sequencing, and then crossed with appropriate balancer stocks. The resulting Pex5.FLAG knock-in flies were maintained as stable homozygous stocks.

### RNA extraction and quantitative RT-PCR

7–10 female flies were homogenized using TissueLyser II (QIAGEN), and RNA was extracted using TRIzol (Thermo Fisher Scientific). DNase-treated total RNA was then reverse transcribed to cDNA using iScript cDNA Synthesis Kit (Bio-Rad). Quantitative RT-PCR (qRT-PCR) was performed with a Quantstudio 3 Real-Time PCR system and PowerUp SYBR Green Master Mix (Thermo Fisher Scientific). Three independent biological replicates were performed with two technical repeats. The mRNA abundance of each candidate gene was normalized to the expression of RpL32 by the comparative CT methods. Primer sequences are listed in the following: RpL32: forward 5’-AAGAAGCGCACCAAGCACTTCATC-3′ and reverse 5’-TCTGTTGTCGATACCCTTGGGCTT-3′; *P*ex5 Primer 1: forward 5′-TGTGCCTGCATGTGGATCAT −3′ and reverse 5′-TCCAGGAGATGCGGGACATA −3′; *P*ex5 Primer 2: forward 5′- CAACCTTACACACCCACATGAC-3′ and reverse 5′-GCAGCGATCTCCAGAGTTAT-3′.

### Adult Brain Immunohistochemistry

Adult female *Drosophila* (7 days old) were anesthetized on ice. Using Dumont 3C tweezers, brain dissections were immersed and dissected in *Drosophila* Schneider’s Media. Brain tissues were immediately fixed in 4% paraformaldehyde (PFA) and kept in a rotator for 25 minutes at room temperature. Subsequently, brains were washed three times for 15 minutes each in 0.2% PBS-TritonX and subsequently kept in blocking solution of 5% normal donkey serum for 1 hour. Tissues were then incubated with primary antibodies for 48 hours overnight at 4°C, followed by three 15 minutes washes of 0.2%PBS-TritonX and then secondary antibody staining for another 48 hours at 4°C. Samples were washed three times for 15 minutes each in 0.2% PBST and then mounted with microfrost slides in Prolong Diamond (Life technologies). Primary antibodies used were as follows: anti-PMP70 (1:200; a gift from Kyu-Sun Lee Lab); anti-ELAV (1:200; DSHB); anti-FLAG(1:200; Thermo); anti-SKL (1:200; a gift from Richard Rachubinski); and anti-nc82 (1:20; DSHB). Secondary antibodies were stained at 1:200 and included 488 donkey anti-mouse, 647 donkey anti-mouse, and 488 anti-guinea Pig (Jackson ImmunoResearch Laboratories Inc, West Grove PA).

### Neutral lipid BODIPY 493/503 staining protocol

A 10mM main stock of BODIPY 493/503 was prepared from 10mg of BODIPY 493/503 by adding 5.2mL of DMSO and mixing until l completely dissolved. Working stocks of 2mM were prepared in DMSO and then used 1:100 (final concentration: 20μM) on samples already fixed and stained for primaries and secondaries (see above). Samples were stained with BODIPY 493/503 (20μM) for 10 minutes, before mounting in glycerol and imaging on an Olympus FV3000 confocal as noted above. For BODIPY experiments, ZT2 was exclusively used.

### Confocal Microscopy

Images were captured using an Olympus FV3000 laser scanning confocal (Olympus, Waltham, MA, USA). 60x and 100x oil-immersive lens were used to capture brains, with further 5x digital zooms into a consistent cortical region, below the antennal lobes and above the suboesophageal ganglion. 10 slices were taken through the cortex. For the circadian import experiments, acquisition of samples was done blind to genotype by independent investigators in two experimental runs. Multiple ROIs (3-5) of 200 x 200 pixel dimensions were acquired per biological replicate brain.

### Image analysis

For Pearson analysis, the *JACoP* plugin tool (*FIJI*) was used to automatically calculate the Pearson correlation coefficient (scale from 0 to 1, with 1 being the highest level of colocalization) between two 16-bit images in different color gradient [111]. For peroxisomal import quantification (SKL and PTS1.YFP), puncta were counted via the cell counter function on FIJI and by blinded independent investigators for 3-5 ROIs of a consistent 200×200 pixel dimension.

For BODIPY lipid droplet analysis, Fiji particle analysis was used to measure lipid droplet size through an automated workflow analysis. Measurements of all replicates of control versus mutant size were plotted in a size difference graph. For lipid droplet total count, multiple ROIs (3-5) of 200 x 200 pixel dimensions were acquired per biological replicate brain to aggregate representative average lipid droplet count of control versus mutant brain samples.

### Pex5.FLAG Quantification

Flies of 7 days old age at various timepoints (ZT2/ ZT8/ZT14/ZT20; CT2/CT8) were dissected and fixed (see above, immunohistochemistry). Imaging was conducted on an Olympus FV3000 confocal. Anterior region of the cortex glial cells and surrounding soma below the antennal lobes and above the suboesophageal ganglion were captured using a 60x oil lens (with 5x digital zoom), with 10 slices taken through the cortex of anterior region of brain. 8-bit image were used for analysis by performing thresholding to highlight representative intensity of FLAG staining compared to anti-ELAV stain of the surrounding soma regions. Measurements were taken of intensity to highlight differences in intensity of FLAG regions versus surrounding soma.

### Lipidomics extraction and LC-MS/MS

20 female brains at day 7 ZT10-12 were dissected rapidly on ice-cold 1X phosphate-buffer saline and frozen on dry ice in chilled methanol (90% v/v) before storage at −80°C until lipid extraction. A modified folch extraction was used and separate columns and standards were run on an Agilent 6430 Triple Quadrupole. Internal standards used included Equisplash™ LIPIDOMIX® Quantitative Mass Spec Internal Standard (Avanti #330731) and Oleic Acid-d9 (Avanti #861809), following a protocol similar to [112].

For free fatty acid analysis, an Agilent Zorbax RX-Sil column 3.0 × 100 mm, 1.8 µm particle size was used. The total LC run time was 19 min (with 2.5 min post-run) at a flow rate of 0.3 µL/min. Mobile phase A was composed of isopropanol/hexane/water (58:40:2, v/v) with 5mM ammonium acetate and 0.1% acetic acid. Mobile phase B consisted of isopropanol/hexane/water (50:40:10, v/v) with 5mM ammonium acetate and 0.1% acetic acid. Gradient elution consisted of 0 to 17 min of 0% mobile phase B and an increase from 17 to 18 min until 100% of mobile phase B and after 18 min the mobile phase decreased to 0% for 1 more min. The Agilent 6430 Triple-Quad LC/MS system (Santa Clara, CA) was operated in negative mode. Free fatty acids were identified as [M - H]^-^ and it was run as a pseudoMRM.

Triglycerides were separate as described by Kehelpannala et al., 2020[113]. An InfinityLab Poroshell 120 EC-C18 2.1 × 100 mm (2.7-Micron particle size) column (Agilent, USA) operated at 55 °C using an Agilent 1260 Infinity LC system (Santa Clara, CA). The solvent system consisted of Mobile phase A acetonitrile (ACN)-water (60:40, v/v) and mobile phase B and isopropanol (IPP)-ACN (90/10, v/v) both containing 10 mM ammonium formate. The gradient was starting with a flow rate of 0.26 mL/min and it was set to first a 0–1.5 min isocratic elution with 32% B which was then increased to 45% B from 1.5 to 4 min, then to 52% B from 4 to 5 min followed by an increase to 58% B from 5 to 8 min. Next, the gradient was increased to 66% B from 8 to 11 min followed by an increase to 70% B from 11 to 14 min and an increase to 75% B from 14 to 18 min. Then, from 18 to 21 min B was increased to 97% and B was maintained at 97% from 21 to 25 min. Finally, solvent B was decreased to 32% from 25 to 25.10 min, and B was maintained at 32% for another 4.9 min for column reequilibration. The Agilent 6430 Triple-Quad LC/MS system (Santa Clara, CA) was operated in positive mode. Triglycerides were identified as [M + NH4]^+^ molecular ions representing the neutral loss of the fatty acid. The collision voltage was 21 eV.

Gas-phase ions of various lipid species were obtained using electrospray ionization (ESI), with the electrospray needle held at −3,500 V (+) and 3,000 V (–). Data acquisition and analysis was carried out using Agilent Mass Hunter software.

### Lipidomic Analysis

Data were analyzed in R studio. Lipids were filtered for abundance across two experimental runs (4 tubes of 20 brains per condition, totaling for controls 8 tubes of *pex5^KD^* control, 4 tubes of Dcr-2;*repo,* 4 tubes of *Dcr2; nSyb*; and for mutants, 8 tubes of pan-glial *pex5^KD^*, 8 tubes of neuronal *pex5^KD^*, and 7 tubes of *CG>pex5^KD^*). Brains were dissected at ZT10-12. For volcano plots, parent control values were averaged (ie *repo-GAL4* and *pex5^KD^* were averaged and compared to *repo-GAL4 > pex5^KD^*). Principal Component Analysis was conducted on sphingolipids, phospholipids, TAG, and free fatty acids using relative % matrices, where the entire lipid composition of a class summed to 100%. Specific ng/brain species plots were analyzed by ANOVA, with Dunnett’s test against pex5^KD^ controls, and Benjamini-Hochberg corrected for further genotypic comparisons. For heatmaps, relative abundance was Z-scored per experimental batch and then pooled.

### Sleep Behavioral Assay

For sleep behavioral analysis in light-dark conditions (12hr light/12-hr dark), 5-day old post eclosion flies were monitored using *Drosophila* Activity Monitors (DAMs, Trikinetics). Male flies were collected at day 5 of adulthood and loaded into DAMs, then analyzed from day 6 to day 10 after a 24 hour recovery period. Cessation of activity for 5 mins was used as a proxy for sleep; flies that were inactive for 12h prior to endpoint of the analyzed period were omitted from analysis as dead. For sleep and activity analysis, code was adopted from[112] and analyzed and graphed in R Studio in a batch function agnostic to genotype. Data were also analyzed using ClockLab and Shiny R-DAM (https://karolcichewicz.shinyapps.io/shinyr-dam/). The Mann-Whitney U test was used to test for significant population differences between controls and mutants.

For free-running rhythmicity assay to assess circadian rhythms and endogenous circadian control of peroxisomal import, five day old flies previously entrained in LD were covered with silver aluminum foil and an opaque black cardboard (DD). Immunohistochemistry on DD brains was conducted after 36 hours of DD, while sleep behavior was collected from day 6 to day 10.

### Heat Map Hierarchical Clustering & Generation

Code was generated by Anurag Das on PositCloud which replicates JUPYTER notebook environment. Code can be shared upon request. After hierarchical clustering, data were further aggregated and visualized using GraphPad Prism.

## QUANTIFICATION AND STATISTICAL ANALYSIS

GraphPad Prism (GraphPad Software Version 10) was used for statistical analysis. Unpaired two-tailed Student’s t- test for comparison between 2 groups or one-way ANOVA (Tukey multiple comparison) was performed to compare the mean value between control and treatment groups of more than two groups respectively. Lipidomics data were analyzed and plotted in R studio.

